# Redirection of SARS-CoV-2 to Phagocytes by Intranasal sACE2-Fc as a Universal Decoy Confers Complete Prophylactic Protection

**DOI:** 10.1101/2025.09.23.677998

**Authors:** Jingyi Wang, Jiangchuan Li, Alex W. Chin, Bin Luo, Junkang Wei, Jiale Qiu, Jianwei Ren, Yin Xia, Thomas Braun, Leo L.M. Poon, Bo Feng

## Abstract

The rapid evolution of SARS-CoV-2 and other respiratory RNA viruses limits the success of current vaccines and antibody-based therapies. Engineered decoy receptors based on soluble angiotensin-converting enzyme 2 (sACE2) offer promising alternatives. Clinical-grade recombinant sACE2 inhibits SARS-CoV-2 replication *in vitro* but shows limited clinical success. This study reports an optimized sACE2 mutant fused to human IgG1 Fc (B5-D3), which redirects virus–decoy complexes to lysosomal degradation in macrophages. Intranasal prophylactic delivery of B5-D3 confers complete protection in SARS-CoV-2-infected K18-hACE2 mice. Abrogation of Fc effector functions compromises antiviral protection, indicating that Fc-mediated uptake of virus–decoy complexes is critical. Transcriptomic analysis suggests that B5-D3 induces early immune activation in lungs of infected mice. Bio-distribution and flow cytometry reveal selective targeting of airway phagocytes. *In vitro* assays confirm lysosomal degradation of virus–decoy complexes by macrophages without productive infection. These findings reveal a distinct antiviral mechanism via phagocytic clearance, supporting refined regimens for decoy treatments against SARS-CoV-2 and potentially other respiratory viruses.

## INTRODUCTION

The incessant evolution of severe acute respiratory syndrome coronavirus 2 (SARS-CoV-2) and frequent breakthrough infections during the coronavirus disease 2019 (COVID-19) pandemic underscore the critical need for effective antiviral strategies that are less susceptible to immune escape than conventional vaccines and monoclonal antibody (mAb) therapies [1].

Soluble angiotensin-converting enzyme 2 (sACE2) therapies, which employ recombinant forms of the human angiotensin-converting enzyme 2 (ACE2) receptor—the primary binding site for the SARS-CoV-2 spike protein [1–5]—as viral decoys, have emerged as a promising alternative [6]. However, an early clinical version (amino acid [aa] 1-740, APN01) showed limited therapeutic benefit [7] and raised safety concerns about interference with endogenous renin-angiotensin system (RAS) [8]. Subsequent protein engineering greatly improved the pharmacological properties of sACE2, including fusion with a human IgG1 Fc domain (sACE2-Fc) to enhance serum half-life [9], and mutagenesis to enhance spike-binding affinity [10–12] and abolish enzymatic activity [10, 12, 13]. Potent sACE2-Fc mutants have shown broad-spectrum neutralization against SARS-CoV-2 variants in animal models [14–16]. However, their efficacy in protecting hosts from viral infections was often incomplete. Despite the evidence suggesting a role for Fc-mediated effector functions in sACE2-Fc efficacies [14], underlying immune mechanisms remained poorly understood. Further investigation that systematically assesses the potential to optimize decoy design, strategies of administration, and mechanisms of actions is pivotal to the development of ACE2 decoy-based antivirals and harnessing their full potential.

In this study, we engineered a potent yet minimally mutated sACE2-Fc decoy candidate (B5-D3) with just two mutations to enhance spike-binding (T92Q) while eliminating enzymatic activity (H374N). Broad-spectrum neutralization capacity against multiple SARS-CoV-2 variants was confirmed by *in vitro* neutralization assays. Markedly, stepwise examinations of various administration routes and time points identified intranasal (IN) prophylaxis as the most effective regimen for B5-D3, which conferred complete protection against SARS-CoV-2 infection in K18-hACE2 mice across age groups. Whereas B5-D3 intravenously administered either prior or post infection, showed activity that moderately improved disease outcome. To understand how sACE2-Fc decoys influence viral fate and achieve superior antiviral protection via IN prophylaxis, we carried out systematic, mechanistic investigations through transcriptomics, bio-distribution, and phagocytosis analysis. Our results revealed that IN-delivered B5-D3 engages airway phagocytes to promote early viral clearance and host immune activation, which uncovers a distinct antiviral mechanism and offers a universal and commonly applicable “decoy strategy” to combat unknown air-borne respiratory virus in the future.

## RESULTS

### Engineered sACE2-Fc decoys with two single mutations achieve robust neutralization against SARS-CoV-2 variants

To generate representative ACE2 decoys with optimal performance, we adopted the established sACE2-Fc fusion design [9] (Fig. 1a; Supplementary Fig. 1) and selectively verified mutation(s) proposed in prior studies, either to enhance the binding of human ACE2 to SARS-CoV-2 spikes [17] (B2–B6) or to abolish enzymatic activity [10–12, 18] (A2, A3, D1–D5) (Supplementary Fig. 2a,b). The sACE2-Fc candidates generated with either type of mutation(s) were all verified for pseudovirus-based neutralization assays [19] and ACE2 enzymatic activity. Indeed, mutants B2–B6 showed consistently enhanced neutralization capacity against both Wuhan-Hu-1 and D614G pseudoviruses [20] (Fig. 1b; Supplementary Fig. 2c,d). Notably, B5 with the single T92Q mutation, which increases spike affinity by removing a critical glycosylation site at N90 [21], exhibited remarkable neutralization enhancement among other multi-mutants. Meanwhile, mutations selected for catalytic inactivation, including the previously reported mutation pairs A2 [11, 18], A3 [10], single mutations derived from these pairs (D1, D3, D4, but not D2), and a recently reported single mutation D5 [12], effectively abolished enzymatic activity while showing minimal effect on spike binding (Fig. 1b; Supplementary Fig. 2e). We next combined the B5 (T92Q) mutation with each of the inactivating mutations. Notably, among the resulting compound mutants, B5-D1, B5-D3, B5-D4, and B5-D5 with two mutations remained to be enzymatically inactive while retaining comparable or stronger neutralization capacity than B5-A2 and B5-A3 with three mutations (Fig. 1b; Supplementary Fig. 2f–h). B5-D3 (T92Q/H374N) emerged as one of the best candidates (Fig. 1a, red stars; Fig. 1b, red arrow), exhibiting minimal deviation from wild type (WT) ACE2 in structural modeling (root mean square deviation [RMSD] = 0.212 Å; Supplementary Fig. 2i) [22].

**Fig. 1.**
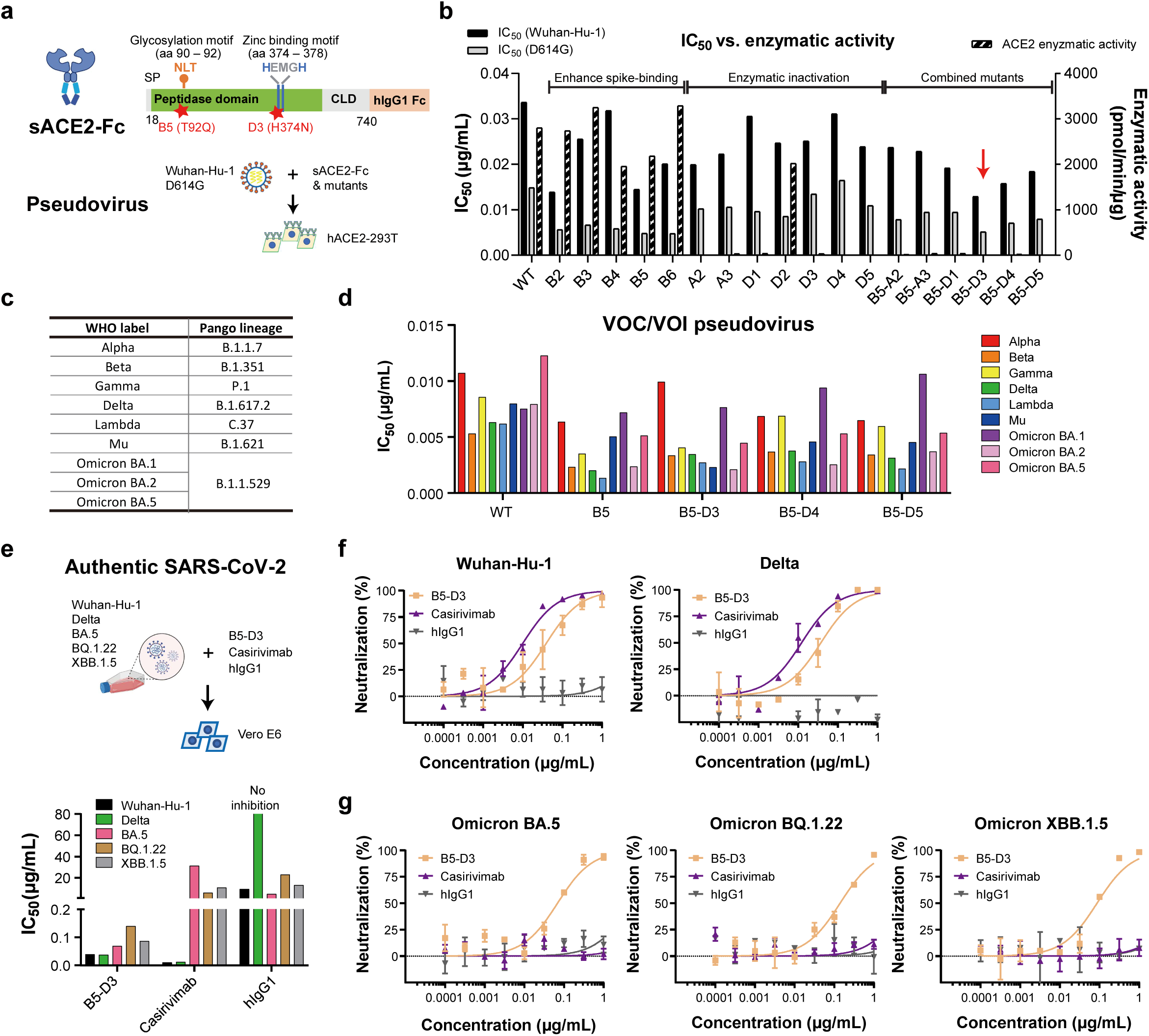
Enhanced sACE2-Fc with two single mutations exhibited broad-spectrum neutralization of SARS-CoV-2 variants. **a** Schematic representation of sACE2-Fc structure (upper) and neutralization assay setup (lower). Key amino acid positions (90-92 and 374-378) involved in glycosylation and zinc binding are highlighted. Red stars mark the positions of mutations in the sACE2-Fc mutant B5-D3. SP, signal peptide; CLD, collectrin-like domain; hIgG1, human IgG1. **b** Comparative bar graph showing the half-maximal inhibitory concentration (IC_50_) values for neutralization of Wuhan-Hu-1 and D614G pseudovirues by WT sACE2-Fc and mutants (B2 to B6, A2, A3, D1 to D5, and B5-derivatives). The red arrow emphasizes the superior performance of the B5-D3 mutant. Enzymatic activity of each construct is plotted on the right axis. **c** List of pseudoviruses carrying spikes from different SARS-CoV-2 variants tested, categorized by the World Health Organization (WHO) into VOCs and VOIs. **d** Graph displaying IC_50_ values of WT sACE2-Fc, B5, and B5-D3/4/5 mutants against various SARS-CoV-2 VOCs and VOIs in neutralization assays. **e** Schematics of the plaque-reduction neutralization tests (PRNTs) process (upper) and the resulting IC_50_ values for B5-D3, Casirivimab, and hIgG1 against authentic SARS-CoV-2 (lower). **f, g** Dose-response curves depicting the neutralization efficacy of B5-D3 (orange), Casirivimab (purple), and hIgG1 (grey) in PRNTs against authentic SARS-CoV-2 Wuhan-Hu-1 and Delta strains (**f**), and Omicron sub-lineages (**g**). Data are presented as mean ± standard deviation (SD) from duplicate experiments.

To assess the breadth of neutralization, we tested three double mutants (B5-D3, B5-D4, and B5-D5) against pseudoviruses bearing spikes from various variants of concern (VOCs) and variants of interest (VOIs) [1, 23–27]. All three designs showed dose-dependent neutralization with higher potency than WT sACE2-Fc (Fig. 1c,d; Supplementary Fig. 3). We further examined B5-D3, as a representative decoy candidate, against authentic SARS-CoV-2 using plaque reduction neutralization tests (PRNTs) in Vero E6 cells, which indeed, confirmed its robust activity against Wuhan-Hu-1, Delta, and Omicron variants BA.5, BQ.1.22, and XBB.1.5 strains [1, 24, 28, 29] (Fig. 1e-g). In contrast, Casirivimab, serving as positive control [30], showed efficacy only against early variants (Wuhan-Hu-1 and Delta; Fig. 1f), but failed to neutralize Omicron sublineages (Fig. 1g). These results demonstrate that a rationally engineered sACE2-Fc decoy with only two mutations could achieve potent and safe neutralization across SARS-CoV-2 variants, reducing the potential risks associated with extensive mutagenesis.

### Prolonged *in vivo* overexpression of sACE2-Fc double mutants demonstrates minimal RAS disturbance and no tissue damage in mice

Next, we evaluated the safety of the sACE2-Fc double mutants *in vivo* (Supplementary Fig. 4a). Adult K18-hACE2 transgenic mice with immune tolerance to human ACE2 [31] were intravenously injected with adenovirus-associated virus (AAV) vectors encoding either WT sACE2-Fc or double mutants at a dose of 1×10^11^ genome copies (GC) per mouse. Notably, serum levels of the double mutants (B5-D3, B5-D4, and B5-D5) were significantly higher than those of WT sACE2-Fc (Supplementary Fig. 4b; Supplementary Table 1). This trend was further supported by quantification of AAV genomes in the liver, indicating greater *in vivo* stability or tolerance of the double mutants (Supplementary Fig. 4c).

Importantly, despite prolonged high-level expression, ELISA measurements of serum renin, Angiotensin II (Ang II), and Ang (1–7) [8] demonstrated minimal disturbance to the RAS in mice treated with the double mutants (Supplementary Fig. 4d–f). In contrast, WT sACE2-Fc treatment led to significantly elevated serum levels of renin and Ang II, indicating a disruption of the RAS (Supplementary Fig. 4d,e). Histological examination of multiple organs at the end point showed no evidence of tissue damage in any of these groups (Supplementary Fig. 4g). These observations collectively underscore the improved safety of catalytically inactive sACE2-Fc mutants, supporting their suitability for prolonged or repeated use.

### Prophylactic administration of the sACE2-Fc B5-D3 mutant via the intranasal route exhibits superior protection against SARS-CoV-2

Next, we evaluated the *in vivo* efficacy of the sACE2-Fc double mutant B5-D3 against SARS-CoV-2 infection using aged K18-hACE2 mice (10 – 12 months old) (Fig. 2a). 6 hours (h) before inoculating with 1 × 10^4^ plaque-forming unit (PFU) of SARS-CoV-2 (Wuhan-Hu-1 strain), mice received a prophylactic dose of recombinant B5-D3 protein either intranasally (IN, 2.5 mg/kg) or intravenously (IV, 15 mg/kg). To simulate a therapeutic intervention, an additional group received IV B5-D3 (15 mg/kg) 24 h post-virus inoculation. The vehicle control group received an intranasal PBS administration 6 h before viral challenge. Over a 14-day observation period, all mice in the PBS group exhibited significant weight loss and succumbed to infection by 7 days post-infection (dpi) (Fig. 2b,c, black lines). Both IV-treated groups exhibited initial weight loss similar to the PBS group; however, two out of four mice in each group began to regain weight from 10 dpi and survived until the observation endpoint, suggesting improvement in disease severity and survival (green and blue lines). Notably, all mice in the IN-prophylaxis group, despite receiving a 6-fold lower dose of B5-D3 protein than those in the IV groups, maintained stable body weight and achieved complete survival over the 14-day period (red lines).

**Fig. 2.**
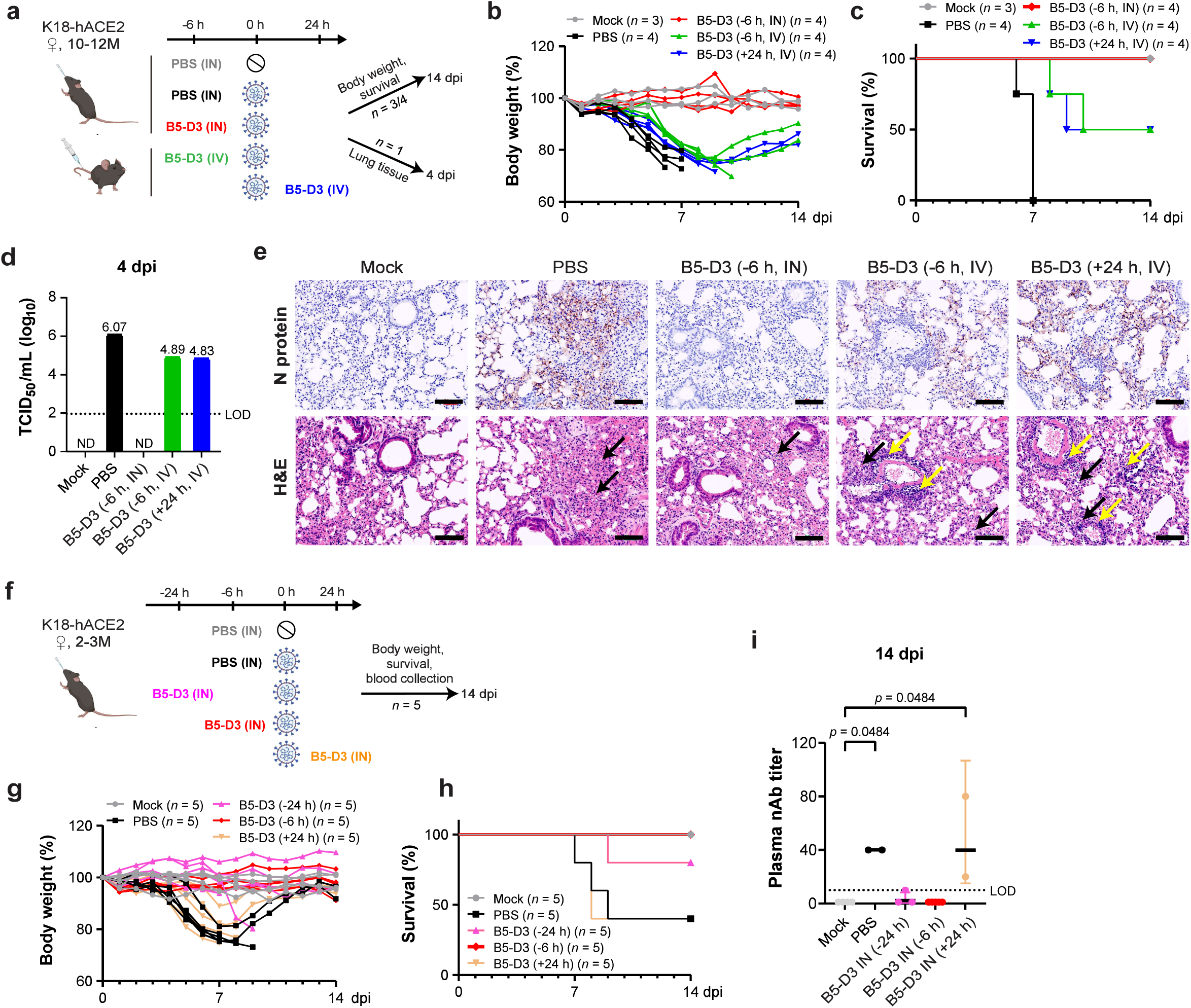
Enhanced survival and reduced infection in K18-hACE2 mice through intranasal prophylaxis with B5-D3 against SARS-CoV-2. **a**–**e** Female K18-hACE2 mice, aged 10 to 12 months, were inoculated with 1 × 10^4^ PFU of SARS-CoV-2 (Wuhan-Hu-1 strain). Mice were treated with B5-D3 6 h prior (–6 h) via intranasal (IN, red) or intravenous (IV, green) routes, or 24 h post-infection (+24 h, blue) via IV (*n* = 4 + 1). IN PBS administered 6 h prior to viral challenge served as the vehicle control (black; *n* = 4 + 1), and PBS alone was used for mock control (grey; *n* = 3 + 1) (**a**). Body weight and survival (*n* = 3 or 4) were monitored over 14 days (**b**, **c**). One mouse from each group was sacrificed at 4 dpi for analysis of viral titers in lung homogenates using a median tissue culture infectious dose (TCID_50_) assay (**d**) and histological analysis of lung sections (upper, IHC staining for N protein; lower, H&E staining) (**e**). Black arrows indicate alveolar thickening, and yellow arrows show leukocyte infiltration. Scale bar = 100 μm. ND, not detected; LOD, limit of detection. **f**–**i** Young female K18-hACE2 mice, aged 2 to 3 months, were inoculated similarly and treated with B5-D3 via IN route at 24 h before (–24 h, pink), 6 h before (–6 h, red), or 24 h after (+24 h, orange) the viral challenge (*n* = 5). Mice receiving IN PBS 6 h before infection served as the vehicle control (black), with mock control mice receiving PBS alone (grey) (**f**). Body weight (**g**) and survival (**h**) were recorded for 14 days. Neutralizing antibody titers against Wuhan-Hu-1 in serum samples from surviving mice at 14 dpi were determined using Vero E6 cells (**i**). nAb, neutralizing antibody. Data are presented as the geometric mean ± geometric SD. Statistical significance was determined using Dunn’s multiple comparisons test.

To monitor viral burden, one mouse from each group was sacrificed at 4 dpi (Fig. 2a). Corroborating the body weight and survival data, no infectious viral particles were detected in lung homogenate from the IN-prophylaxis mouse. In contrast, mice treated with IV prophylaxis or therapy showed reduced but still detectable viral titers compared to the PBS group (Fig. 2d). Immunohistochemistry (IHC) staining further confirmed the absence of viral nucleocapsid (N) protein in the IN-treated mouse, whereas IV-treated mice showed residual infection and immune cell infiltration. H&E staining revealed varying degrees of alveolar thickening in all SARS-CoV-2-inoculated mice (Fig. 2e).

To further explore the effective timing of IN administration that offers superior antiviral protection, we treated a younger cohort of K18-hACE2 mice (2 – 3 months old) with B5-D3 (IN, 2.5 mg/kg) at –24 h, –6 h, or +24 h relative to viral challenge (Fig. 2f). Consistently, all mice in the PBS group of the young cohort exhibited substantial weight reduction from 4 dpi and reached approximately 20% loss by 7 dpi (Fig. 2g, black lines). Whereas differently from the aged cohort, two of the five infected young mice eventually recovered, resulting in 40% survival (Fig. 2g,h, black lines) indicating age-related fitness [32]. Interestingly, both the –24 h and –6 h IN-prophylaxis groups maintained stable body weights (Fig. 2g, pink and red lines), ending up with survival rates of 80% and 100%, respectively (Fig. 2h). In contrast, the +24 h IN-therapy group showed substantial weight loss and no survival improvement compared to the PBS group, indicating that high-efficiency antiviral protection of IN B5-D3 is limited to prophylactic administration but not post-infection treatments (Fig. 2g,h, orange lines). Consistently, virus-neutralizing antibodies were detected in surviving mice from the PBS and +24 h groups at 14 dpi, indicating once active infection and subsequent immune response. Whereas antibody levels remained minimal in the two IN-prophylaxis groups, suggesting effective prevention of viral replication (Fig. 2i).

### Efficient protection against SARS-CoV-2 by intranasal B5-D3 prophylaxis depends on Fc-mediated effector functions

Adding on to the 2-week observation of the significant protection conferred by IN prophylaxis with B5-D3, we examined the early responses following SARS-CoV-2 challenge in K18-hACE2 mice. A new cohort of 2- to 3-month-old mice received B5-D3 IN treatment 6 h before infection (–6 h), and lung tissues were harvested at 1, 2, and 4 dpi for analysis (Fig. 3a). An additional group was treated with a modified version of B5-D3, which contains L234A/L235A mutations in the human IgG1 Fc region (B5-D3-LALA) to abolish its binding to Fc gamma receptor (FcγR) and abrogate Fc effector functions [33] (Supplementary Fig. 5). Quantitative PCR of viral spike (*S*) and nucleocapsid (*N*) RNA in lung tissues revealed only marginal viral loads in the B5-D3-treated mice at as early as 1 dpi, indicating efficient suppression of early viral replication compared to the PBS group (Fig. 3b). Analysis of infectious viral particles in lung homogenates further corroborated these observations, demonstrating minimal or undetectable viral titers in the B5-D3 group at all time points (Fig. 3c). In contrast, PBS-treated mice exhibited consistently high viral loads. Interestingly, the analysis of B5-D3-LALA group detected varied levels of *S* and *N* RNAs and significant viral burdens in two out of three mice, indicating partial protection and supporting that Fc effector functions are indispensable for full efficacy of IN B5-D3 (Fig. 3b,c, right panels). Consistently, IHC staining for N protein in lung sections confirmed the absence of viral infection in the B5-D3 groups at all time points. Whereas signs of viral replication were evident in the lungs of mice treated with PBS since as early as 1 dpi and in the B5-D3-LALA-treated cohort as examined at 4 dpi (Fig. 3d; Supplementary Fig. 6, left panels). Despite variations in viral burden, H&E staining indicated alveolar septal thickening in all groups (Fig. 3e; Supplementary Fig. 6, right panels; Supplementary Fig. 7). Notably, moderate alveolar thickening persisted in the B5-D3-treated mice till the end point at 4 dpi, whereas the PBS groups developed much severer alveolar thickening at 4 dpi. Consistent with the partial protection observed with B5-D3-LALA, histological analysis of lung samples in this group revealed severer yet heterogenous alveolar thickening (Supplementary Fig. 7). These findings collectively demonstrated that IN prophylaxis with B5-D3 blocks SARS-CoV-2 infection not only by neutralization but also by immune mechanisms such as Fc-mediated effector functions.

**Fig. 3.**
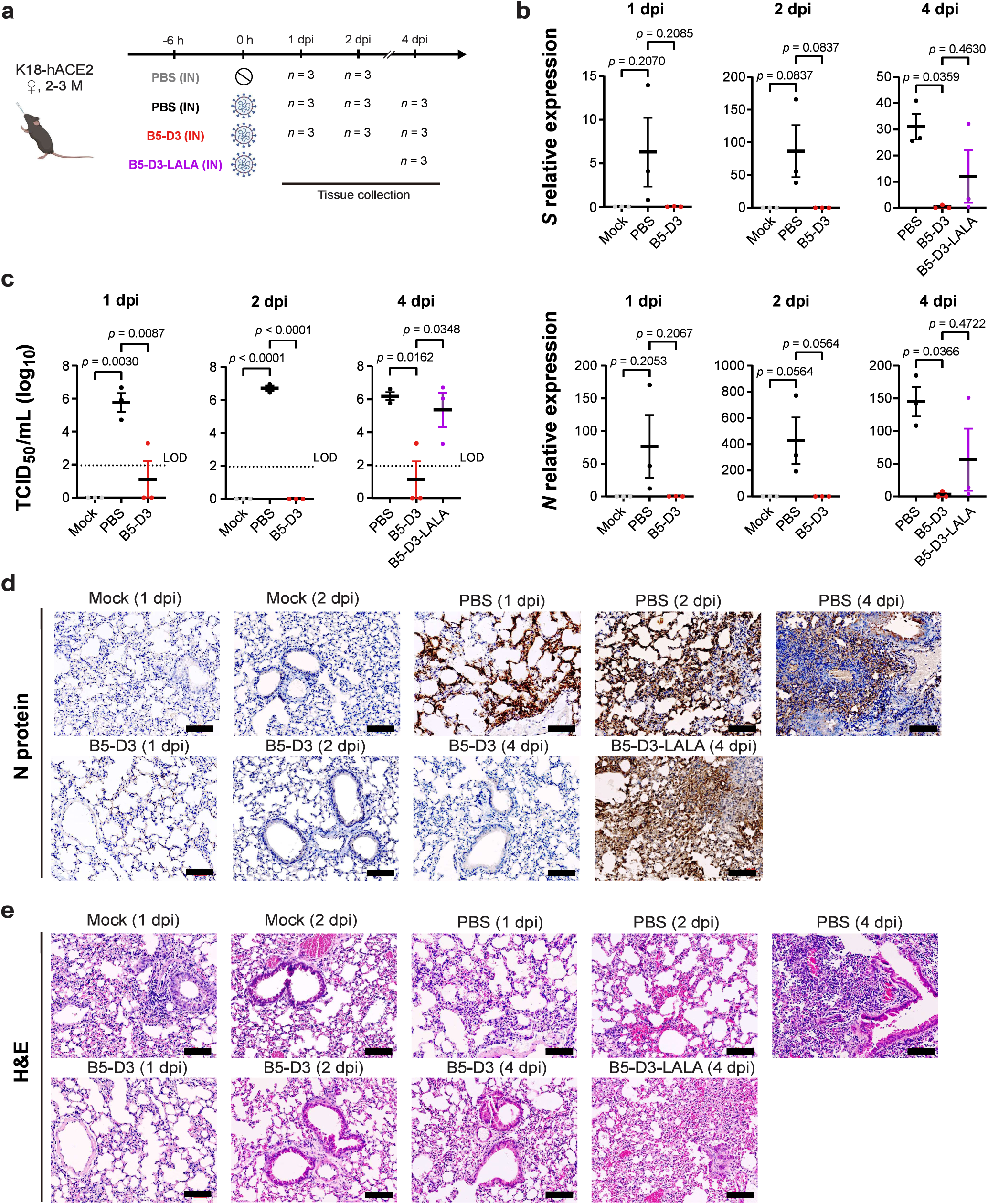
Efficient viral clearance at early stages through intranasal prophylaxis with B5-D3 against SARS-CoV-2 challenge in K18-hACE2 mice. **a** Workflow diagram showing timelines and treatments for different mouse groups. Young female K18-hACE2 mice aged 2 to 3 months received prophylactic administration of PBS (black), B5-D3 (red), or B5-D3-LALA (purple) via the IN route 6 h prior to inoculation with 1 × 10^4^ PFU of Wuhan-Hu-1. Mice inoculated with PBS instead of the virus served as mock controls (grey). Mice from each treatment group were sacrificed for tissue collection at 1, 2, and 4 dpi (*n* = 3 per time point). **b** Quantitative PCR results showing relative amounts of *S* (upper) and *N* (lower) viral RNA in lung tissues collected from different groups at 1, 2, and 4 dpi, normalized to mouse *Gapdh*. **c** The titers of infectious viruses detected in lung homogenates, measured by TCID_50_ assays at 1, 2, and 4 dpi. **d, e** Fixed lung tissues were sectioned and stained; IHC for viral N protein (**d**) and H&E staining for tissue damage (**e**) are shown (scale bar = 100 μm). Data presented as mean ± standard error of the mean (SEM). Statistical significance was determined by Tukey’s multiple comparisons test.

### RNA-Seq analysis of lung transcriptomes reveals early antigen presentation and prompt viral clearance following SARS-CoV-2 neutralization by B5-D3

To delineate the immune mechanisms underlying IN B5-D3-mediated prophylactic protection against SARS-CoV-2, we examined the transcriptomes of lung samples collected at 1, 2, and 4 dpi from the above experiment (Fig. 4a–d; Supplementary Fig. 8a). Unsupervised clustering based on Pearson correlation distinguished samples with severe infection (mainly PBS-treated) from those with subtle or no infection (mocks and most decoy-treated mice) (Supplementary Fig. 8a). Corroborating the levels of viral infections observed, differential gene expression (DGE) analysis revealed extensive inflammatory responses in the PBS groups, significantly greater than in mock treatments. At 1, 2, and 4 dpi, 26, 1232, and 1756 genes were upregulated, respectively, and were significantly enriched in Gene Ontology Biological Process (GOBP) terms related to antiviral responses such as type I interferon (IFN) responses and innate immune responses (Fig. 4a,b; Supplementary Fig. 8b–d) [34]. In stark contrast, DGE analysis between B5-D3 prophylaxis and mocks at 1, 2, and 4 dpi showed subtle changes, with only 1, 7, and 32 genes upregulated, respectively, and only moderate enrichment in chemotaxis-related pathways at 4 dpi (Fig. 4c; Supplementary Fig. 8e). The B5-D3-LALA group, however, had 264 genes upregulated at 4 dpi compared to the mocks, suggesting incomplete protection and ongoing viral activity (Fig. 4d; Supplementary Fig. 8f).

**Fig. 4.**
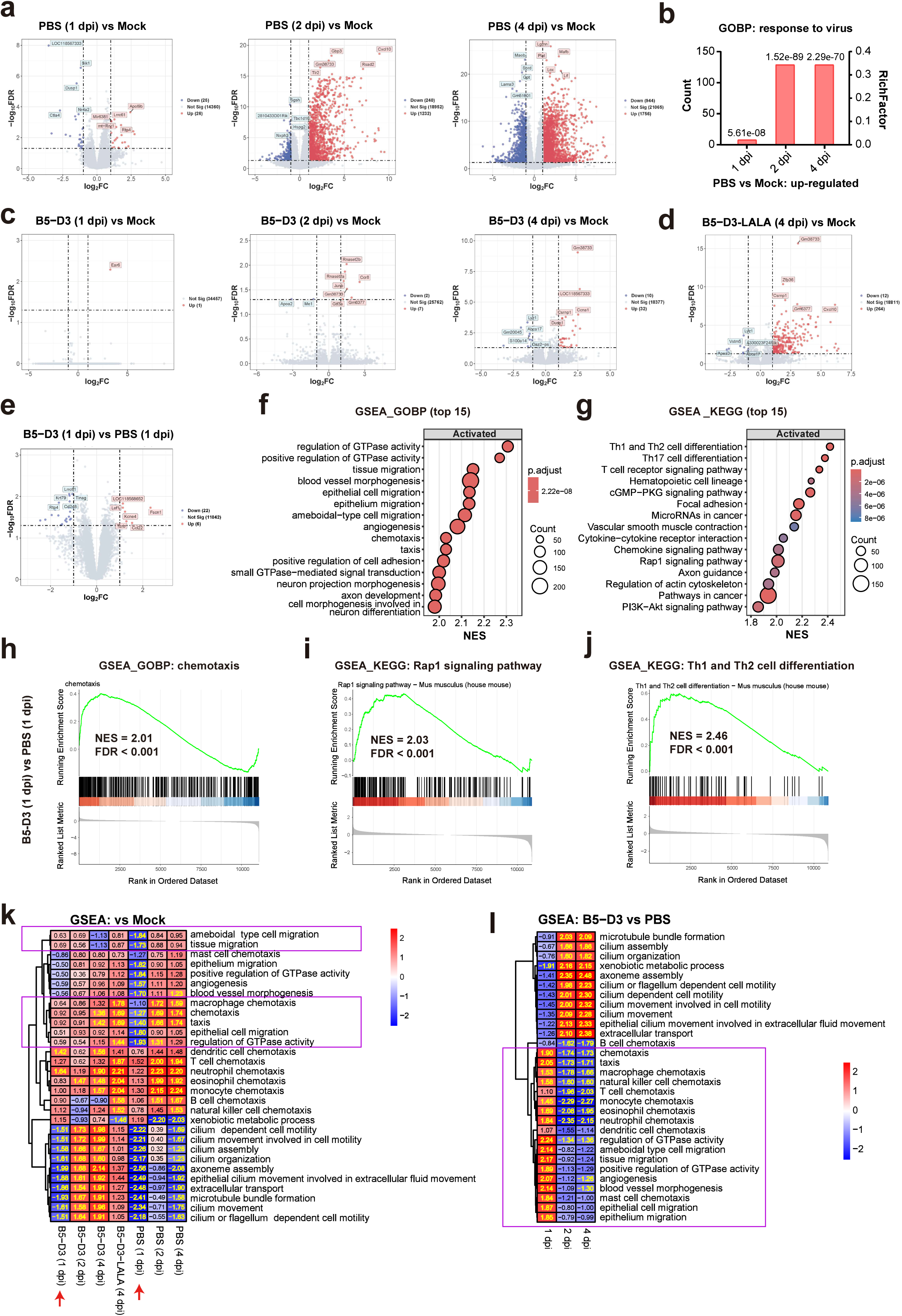
Transcriptomic analysis of lungs revealed early immune activation in IN B5-D3-prophylaxis mouse group after SARS-CoV-2 challenge. **a**–**d** DGE analysis comparing PBS (**a**), B5-D3 (**c**), and B5-D3-LALA (**d**) against the mock control at specific time points (*n* = 3). Volcano plots illustrate the gene expression changes (**a**, **c**, **d**), while red and blue dots represent significantly upregulated and downregulated genes, respectively, with |log_2_ fold change (log_2_FC)| ≥ 1 and a false discovery rate (FDR) < 0.05. Bar chart in **b** shows the enrichment of GOBP “response to virus” observed in PBS groups at 1, 2, 4 dpi, in which adjusted *p* values are indicated for individual comparisons. **e**–**g** Comparison between IN B5-D3 and PBS group at 1 dpi. Volcano plot illustrates the DGE analysis between IN B5-D3 to PBS group at 1 dpi (**e**), with red and blue dots representing significantly upregulated and downregulated genes, respectively, with |log_2_FC| ≥ 1 and FDR < 0.05. GSEA shows top 15 significantly activated GOBPs (**f**) and KEGG pathways (**g**) in IN B5-D3 compared to PBS group at 1 dpi. NES, normalized enrichment score; p.adj, adjusted *p* value. **h**–**j** GSEA plots of chemotaxis (**h**), Rap1 signaling pathway (**i**), and Th1 and Th2 cell differentiation (**j**) in B5-D3 vs PBS comparison at 1 dpi. **k**, **l** Heatmaps show NES of GSEA comparing various treatments to the mock control (**k**) and between B5-D3 to PBS (**l**), focusing on top 10 GOBPs in **f** and **Supplementary Fig. 10c, d**, respectively, and those related to immune cell chemotaxis. Significant NES values (*p* < 0.05, FDR < 0.25) are highlighted in yellow. Purple boxes indicate GOBPs where B5-D3 (1 dpi) group shows activation but PBS (1 dpi) group shows suppression. Benjamin–Hochberg method was used for FDR adjustment.

To capture the immune activations specifically linked to B5-D3-triggered antiviral efficacy other than infection-induced inflammation, we directly compared the B5-D3 and PBS groups (Fig. 4e-j; Supplementary Fig. 9 and 10). Interestingly, at 1 dpi, the B5-D3 group exhibited enhanced expression of several immune-related genes, including *Lef1* [35], *Fscn1* [36], *Kcne4* [37], *Tcrb*, and *Ccl22* [38, 39], which are associated with early dendritic cell function and T cell activation (Fig. 4e). Gene Set Enrichment Analysis (GSEA) of GOBPs and Kyoto Encyclopedia of Genes and Genomes (KEGG) pathways further supported these findings (Fig. 4f,g). Chemotaxis and pathways related to antigen presentation such as Rap1 signaling pathway [40] and Th1 and Th2 cell differentiation were significantly activated in the B5-D3 group at 1 dpi compared to PBS group (Fig. 4h-j; Supplementary Fig. 9a–c). Moreover, the B5-D3 groups showed enhancement in cilium movement and metabolism of xenobiotics at both 2 and 4 dpi, suggesting active clearance of viral particles due to effective early responses (Supplementary Fig. 10c–f).

Furthermore, we collectively examined the GOBPs that were significantly activated in B5-D3 groups at either 1, 2, or 4 dpi among all treatment groups and time points. Markedly, B5-D3 group showed higher normalized enrichment scores (NES) in chemotaxis-related GOBP pathways than PBS group at 1 dpi (Fig. 4k, purple boxes), while direct comparison between B5-D3 and PBS groups further revealed the broad involvement of multiple types of effector immune cells (Fig. 4l, purple box). These results collectively indicate that early immune activation is a hallmark of B5-D3-mediated protection.

Finally, the lung transcriptomes from mice receiving B5-D3 without viral inoculation showed high similarity to the PBS vehicle controls (Supplementary Fig. 11a). The 10 upregulated genes identified showed poor correlation with the virus-inoculated B5-D3 group (Supplementary Fig. 11b,c), supporting that early immune responses observed in B5-D3 IN prophylaxis groups were primarily triggered by virus neutralization rather than by B5-D3 alone.

### Intranasally delivered B5-D3 is enriched in the respiratory tract and targets mainly the airway macrophages

The superior prophylactic antiviral effects of IN over IV administration of B5-D3 as observed in the K18-hACE2 infection experiments (Fig. 2a–e) suggested the importance of mobilizing the local immunity within the respiratory tract. Next, we labeled B5-D3 protein with Alexa Fluor 750 (AF750) and examined its bio-distribution and kinetics after IN administration (Fig. 5a). *In vivo* imaging showed that fluorescence-labeled B5-D3 (B5-D3-AF750) was present in the nasal cavities for at least 24 h after a single IN dose in K18-hACE2 mice (Fig. 5b). *Ex vivo* images further revealed that B5-D3-AF750 distributed in the respiratory tract from nasal cavity to lung within 20 min and remained enriched in lungs by 24 h after administration (Fig. 5c). In contrast, non-respiratory organs showed minimal signals, which were merely detectable in urinary system and liver (Fig. 5d).

**Fig. 5.**
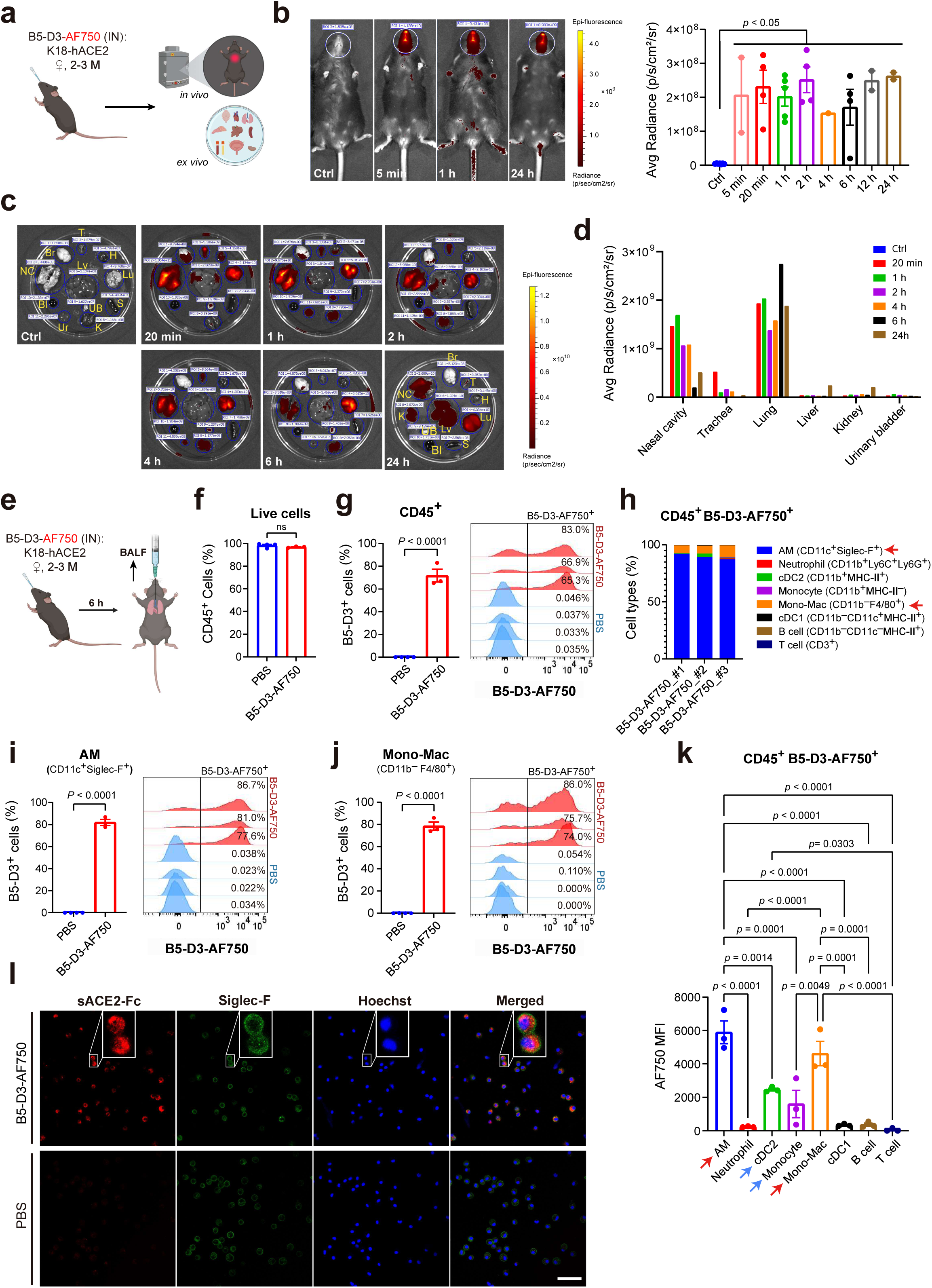
*In vivo* bio-distribution of B5-D3 after IN administration. **a** Schematic workflow of *in vivo* and *ex vivo* imaging. Female K18-hACE2 mice aged 2 to 3 months received IN administration of fluorescently labeled B5-D3 (B5-D3-AF750) and was visualized at different time points. **b** Representative whole-body images of control and treated mice at 5 min, 1 h, and 24 h after B5-D3-AF750 administration, showing the signal captured by *in vivo* imaging (left). White circles indicate regions of interest (ROIs) for quantification of fluorescence signals in the nasal cavities. Average (Avg) Radiance measured at all time points are shown on the right. **c** *Ex vivo* images of tissues from control and treated mice sacrificed at indicated time points after B5-D3-AF750 administration. Blue circles indicate ROIs for signal quantification. Br, brain; NC, nasal cavity; T, trachea; Lu, lung; H, heart; Lv, liver; S, spleen; K, kidney; UB, urinary bladder; Bl, blood; Ur, urine. **d** Avg Radiance shows the fluorescence signals in excised tissues measured *ex vivo*. **e** Schematic workflow for BALF analysis. Female K18-hACE2 mice aged 2 to 3 months received IN administration of B5-D3-AF750 (*n* = 3) or PBS (*n* = 4) and were sacrificed at 6 h later for collection of BALF cells. **f** Percentage of CD45^+^ cells in live BALF cells. **g** Positive rates (left) and histograms (right) of B5-D3 binding/uptake in CD45^+^ BALF cells. Histograms show B5-D3-AF750 fluorescence intensities in CD45^+^ BALF cells from individual mice. **h** Frequency of individual immune cell types in CD45^+^B5-D3^+^ BALF cells. Red arrows point out AMs and mono-Macs with high abundance. AM, alveolar macrophage; Mono-Mac, monocyte-derived macrophage; cDC1/2, type 1 or 2 conventional dendritic cells. **i**, **j** Positive rates (left) and histograms (right) of B5-D3 binding/uptake in CD11c^+^Siglec-F^+^ AMs (**i**) and CD11b^-^F4/80^+^ mono-Macs (**j**). **k** Median fluorescence intensity (MFI) of AF750 indicates B5-D3 binding/uptake in different CD45^+^B5-D3^+^ populations. **l** Confocal images (scale bar = 50 μm) of BALF cells collected at 6 h and stained for sACE2-Fc (red, anti-Fc, Abcam #ab98596), Siglec-F (green, BD #564514), and nuclei (blue, Hoechst). Magnified views are shown in white rectangles. Data are presented as mean ± SEM, and statistical significance was determined by Tukey’s multiple comparisons test or Student’s t-test.

To identify the immune cells in the respiratory tract that are actively engaged with IN B5-D3, we performed flow cytometry analysis on bronchoalveolar lavage fluid (BALF) at 6 h after IN administration of B5-D3-AF750 (Fig. 5e). Like normal conditions, over 95% of live BALF cells were CD45^+^ immune cells, predominantly composed of CD11c^+^Siglec-F^+^ resident alveolar macrophages (AMs) (> 50%) in both treatment and vehicle groups (Fig. 5f; Supplementary Fig. 12). Notably, the IN administered B5-D3-AF750 was actively retained in the CD45^+^ cells, with positive rates exceeding 65% in all treated mice (Fig. 5g). Among the CD45^+^B5-D3^+^ cells, more than 95% were macrophages, composed primarily of CD11c^+^Siglec-F^+^ AMs (87.2 – 91.7%) and Siglec-F^-^CD11b^-^F4/80^+^ monocyte-derived macrophages (mono-Macs; 6.6 – 9.9%) (Fig. 5h, red arrows). Consistently, these macrophage populations also exhibited the highest B5-D3 positive rates (Fig. 5i,j; Supplementary Fig. 13) and greatest median fluorescent intensities (MFI) (Fig. 5k, red arrows) among all immune cell types in the BALF, indicating the strongest B5-D3-AF750 uptake. Other phagocytic cell types such as the type 2 conventional dendritic cells (cDC2) and monocytes also exhibited considerable AF750 intensities (Fig. 5k, blue arrows; Supplementary Fig. 13b,c), suggesting potential relationships between B5-D3 uptake and phagocytic activities. Confocal microscopy of BALF cells after immunostaining further confirmed that the B5-D3-AF750 were present in the cytoplasm after being retained in AMs (Fig. 5l). These results demonstrate that IN B5-D3 preferentially accumulates in the respiratory tract and is predominantly taken up by airway macrophages, supporting their important role in mediating early immune responses.

### sACE2-Fc facilitates phagocytosis of SARS-CoV-2 pseudovirus via mechanisms distinct from ACE2-dependent viral infection

To examine the implication of macrophage involvement in the early immune activation observed in IN B5-D3 treatment groups, we performed cellular analysis using THP-1 cells as an *in vitro* model for phagocytes [41] and examined the sACE2-Fc-dependent phagocytosis of spike-pseudotyped lentiviruses. Indeed, immunostaining of HIV capsid protein p24 confirmed the attachment and entry of pseudoviruses in the THP-1 cells in a B5-D3-dependent manner, with an evidenced signal peak at 6 h post-co-incubation (Supplementary Fig. 14). Interestingly, analysis on the THP-1-derived M0 and M1 macrophages detected even greater p24 signals, indicating stronger phagocytosis activities compared to undifferentiated THP-1 cells (Fig. 6a,b; Supplementary Fig. 15a,b,d,e). This process resembled antibody-dependent cellular phagocytosis (ADCP), which is significant in THP-1-derived M0 and M1 macrophages [42]. Consistently, further examination revealed colocalization of internalized pseudovirus with lysosomal associated membrane protein 1 (LAMP1), indicating trafficking to lysosomes potentially for degradation [43] (Fig. 6c; Supplementary Fig. 15c,f).

**Fig. 6.**
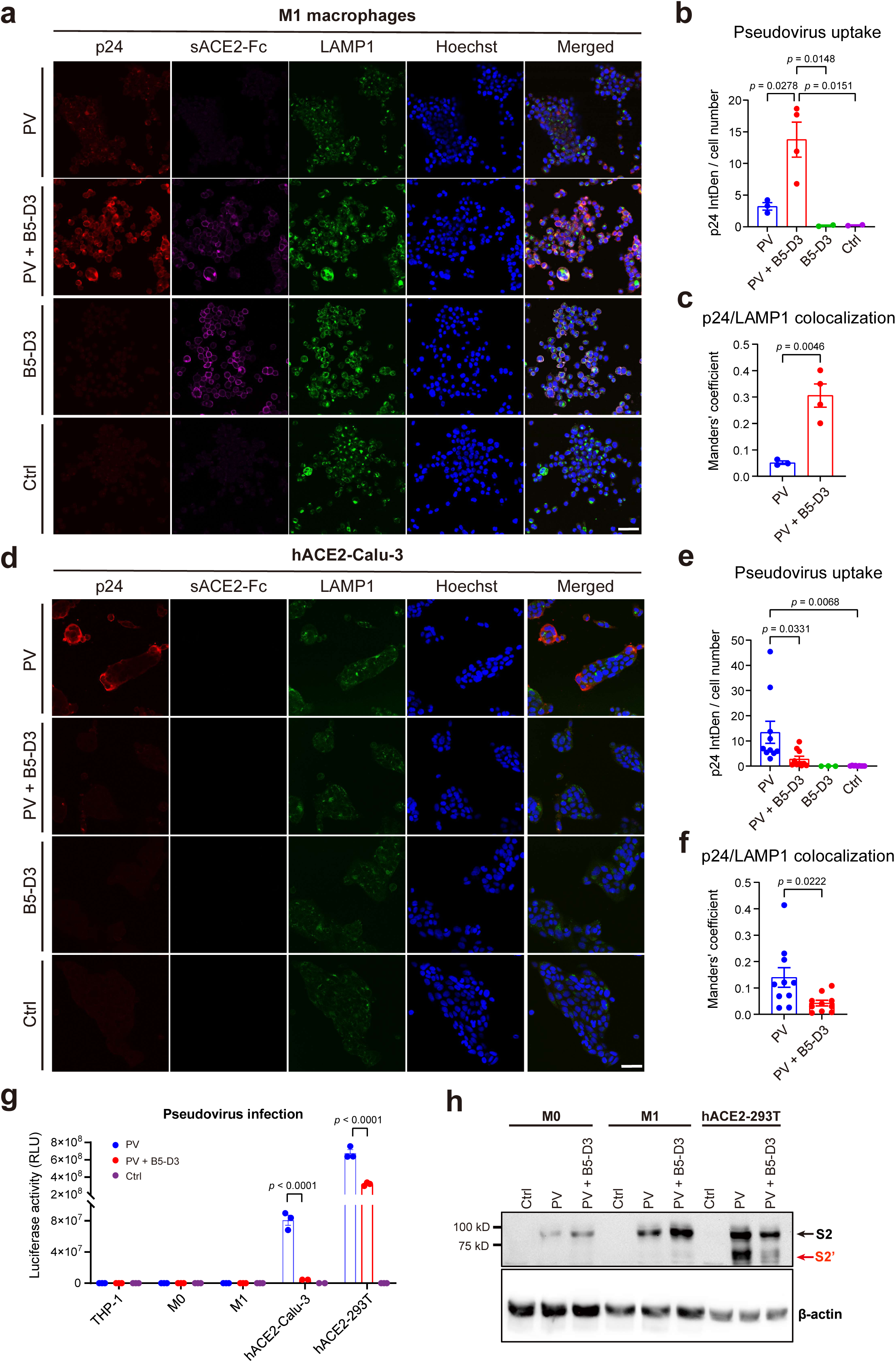
B5-D3 enhanced phagocytosis and degradation of SARS-CoV-2 pseudovirus in THP-1-derived macrophages. **a** Immunostaining of p24 (Invitrogen #PA5-81773), sACE2-Fc, and LAMP1 (Abcam #ab25630) in THP-1-differentiated M0 macrophages showing phagocytosis of SARS-CoV-2 pseudovirus (PV, p24^+^) after 6 h of incubation with or without B5-D3 (scale bar = 50 µm). LAMP1 was stained to identify lysosomes. **b** Quantification of p24 signal intensity as shown in **a**. Intensity Density (IntDen) per cell number indicates the mean p24 signal per cell, calculated using ImageJ. Each dot represents one image. **c** Manders’ coefficient indicating the colocalization of p24 and LAMP1 in THP-1 M0 macrophages as shown in **a**. **d** Immunostaining of p24, sACE2-Fc, and LAMP1 in hACE2-Calu-3 cells after 6 h incubation with pseudovirus, with or without B5-D3 (scale bar = 50 µm). **e** Quantification of mean p24 signal intensity as shown in **d**. **f** Manders’ coefficient for the colocalization of p24 and LAMP1 in hACE2-Calu-3 cells, as shown in **d**. **g** Quantification of pseudovirus infection in THP-1, M0 macrophages, M1 macrophages, hACE2-Calu-3, and hACE2-293T cells, in the presence or absence of B5-D3. Results shown are luciferase activities measured at 2 days post-transduction. **h** Immunoblot staining of cell lysates to detect SARS-CoV-2 spike cleavage after cell entry. M0 macrophages, M1 macrophages, and hACE2-293T cells were incubated with pseudovirus for 6 h, with or without B5-D3, before protein extraction. Band locations of SARS-CoV-2 spike S2 and S2′ fragments are labeled in black and red respectively. Data are presented as mean ± SEM, and statistical significance was determined by Tukey’s multiple comparisons test.

We further examined the pseudovirus uptake in Calu-3 cells overexpressing human ACE2 (hACE2-Calu-3; Supplementary Fig. 16) as a model of lung epithelial cells. In contrast to that in macrophages, co-incubation with B5-D3 significantly reduced the pseudovirus entry in hACE2-Calu-3 cells (Fig. 6d–f). Interestingly, while the pseudovirus transduction in hACE2-Calu-3 cells, in the absence of B5-D3, produced robust luciferase signal indicating viral genome release after cell entry, the evident pseudovirus uptake facilitated by B5-D3 in the THP-1 and derivative macrophages yielded no detectable luciferase activity, which further supported viral degradation within phagolysosomes (Fig. 6g). Corroborating these observations, western blot analysis showed absence of cleaved S2′ fragments in the macrophages that had internalized pseudovirus-B5-D3 complexes, supporting that the pseudoviruses did not undergo membrane fusion or cytosolic release as it did in the epithelial cell model [44] (Fig. 6h).

To evaluate the contribution of Fc-mediated effector functions to virus uptake in macrophages, we further compared the efficiencies of B5-D3, B5-D3-LALA, and hIgG1 isotype in mediating pseudovirus uptake by THP-1-derived macrophages (Supplementary Fig. 17). As indicated by p24 staining, functional impairment (LALA mutations) of Fc in B5-D3 significantly reduced the efficiency of virus uptake, emphasizing the importance of intact Fc in B5-D3 function as decoy. Whereas presence of hIgG1 isotype control showed no impact on pseudovirus uptake, supporting that the phagocytosis was largely specific to the pseudovirus-decoy complexes.

To further evaluate the responses triggered by the pseudovirus-B5-D3 complex in macrophages, we performed RNA-seq analysis. The transcriptomic profiling revealed little difference and identified no DEG between the macrophages incubated with both pseudovirus and B5-D3 and those treated with pseudovirus only (Supplementary Fig. 18a). Whereas interestingly, GSEA detected broad activation of pathways related to antiviral response and macrophage activation in the macrophages internalizing pseudovirus and B5-D3 (Supplementary Fig. 18b–i), corresponding to ADCP effect. These findings indicate moderate immune activation triggered by phagocytosis of pseudovirus-decoy complexes, corroborating with the mild immune activation that accelerated antiviral responses in our mice infection experiments.

Collectively, these findings suggest that IN B5-D3 not only blocks viral entry into epithelial cells but also actively redirects SARS-CoV-2 to phagocytic clearance by engaging airway phagocytes via Fc-dependent mechanisms. Moreover, such ADCP-like process contributes to early immune activation and restricts the infection at the respiratory mucosal surface.

## DISCUSSION

In this study, we comprehensively evaluated the protective efficacy and mechanistic basis of an optimized representative sACE2-Fc decoy (B5-D3) against SARS-CoV-2 infection. By introducing only two mutations (T92Q and H374N), we generated a minimally engineered sACE2-Fc mutant (B5-D3) that achieved broad-spectrum neutralization with minimal risk of disrupting the RAS. Among various administration routes and dosing schedules examined, we demonstrated that IN prophylaxis of B5-D3 achieved the most robust protection, completely preventing disease in both young and aged K18-hACE2 mice. Transcriptomic analysis of the infected lung samples at early time points revealed distinct IN B5-D3-dependent immune activation at the onset of infection, indicating that B5-D3 acted not only as a receptor decoy but also as an immune engager upon viral infection. Bio-distribution analysis of fluorescence-labeled B5-D3 demonstrated rapid uptake and high accumulation in the respiratory tract, primarily within airway macrophages. Phagocytosis assays further supported that sACE2-Fc decoy mediated a rapid viral clearance in macrophages, while abolishing membrane ACE2-mediated infection in epithelial cells. Together, these findings reveal a dual-function mechanism for sACE2-Fc decoys in redirecting SARS-CoV-2 to phagocytic clearance and rapid immune engagement, supporting their potential as intranasal prophylactics against respiratory viruses.

Previous studies have reported multiple ACE2 mutations with remarkable potential in neutralizing SARS-CoV-2. However, in these studies, combining ACE2 mutations based on *in silico* predictions to both enhance spike binding and eliminate the ACE2 enzymatic activity in a single design resulted in accumulation of mutations such as K31F/N33D/H34S/E35Q/H345L [12] and L79F/M82Y/Q325Y/H374A/H378A [14]. These extensive mutations have been implicated in structural instability [12] and reduced production efficiency [17]. More importantly, the high mutation loads raise risks for immunogenicity, which is a critical issue when considering clinical applications. Corroboratively, Urano *et al.* detected *in vitro* T cell stimulation elicited by the L79F mutation, whereas the T92Q mutation showed much lower immunogenicity and enhanced spike binding affinity [45]. In our ACE2 decoy design, we incorporated only two mutations (like T92Q and H374N in B5-D3) to enhance neutralization potency while eliminating enzymatic activity, resulting in simplest compound ACE2 mutants. The B5-D3, as a representative, exhibited not only minimal mutation-related risks but also top-level neutralization potencies among all candidate mutants we tested. Hnece, by coupling structural engineering (Fc fusion for avidity improvement) and mutagenesis (potency and safety optimization), we provided simplest decoy design and further consolidated the generalizable framework for decoy design, which is invaluable to combat a novel, highly contagious, and fatal virus in future, before more effective vaccines and drugs are established.

Previous studies have reported that IN prophylaxis with sACE2-Fc mutants or monoclonal antibodies [15, 16, 46] could effectively protect against SARS-CoV-2, in which the Fc-effector functions were indispensable [14]. However, the precise mechanism has not been well depicted, which prevents the further development of these approaches for translation. Here, our study provided evidence for a deeper mechanistic insight, showing that IN sACE2-Fc decoys rapidly engage host immunity in the respiratory tract. Consistently with others’ work [14], IN delivery of a Fc-null variant (B5-D3-LALA) resulted in suboptimal protection and higher viral infection compared to B5-D3. Notably, despite the minimal infection observed, B5-D3-treated mice showed robust early immune activation, including induction of antigen presentation and T cell activation within 24 h post-infection (Fig 4). These results support that B5-D3 not only neutralizes virus but also primes innate and adaptive immune responses, counteracting early-stage viral immune evasion.

Furthermore, our bio-distribution data showed that IN-delivered B5-D3 preferentially accumulates in the respiratory tract (Fig. 5a–d). Flow cytometry and confocal imaging confirmed strong binding and uptake of B5-D3 by airway phagocytes, primarily alveolar macrophages and monocyte-derived macrophages (Fig. 5e–l). Notably, phagocytosis assays demonstrated that B5-D3–virus complexes were trafficked to lysosomes for degradation in macrophages. Functional impairment in the B5-D3 structure reduced the uptake of virus by macrophages. These findings support a mechanism in which B5-D3 redirects viral particles away from membrane ACE2-dependent epithelial entry and toward phagocytic clearance (Fig. 7). Importantly, RNA-Seq analysis of macrophages treated with virus-decoy complexes suggests that such ADCP-like process also facilitated early immune activation and initiated downstream antiviral signaling cascades before the virus reaches epithelial targets (Fig. 7b). Hence, the IN prophylaxis offers a unique advantage by enabling localized immune priming and efficient viral clearance at the frontline of infection. The IN-administered antibody drugs can share similar mechanism and confer effective protection against primary infection of SARS-CoV-2 or other respiratory viruses. However, antibody drugs can be easily escaped when new viral variants emerge, which is specifically common among rapidly spreading respiratory RNA viruses [47]. In contrast, the decoy strategy is intrinsically resistant to “antigenic escape” of the mutating virus, which is further supported by the trend observed in viral evolution that later-emerging SARS-CoV-2 variants exhibit a higher affinity for the ACE2 receptor, enhancing their infectivity and transmissibility [16].

**Fig. 7.**
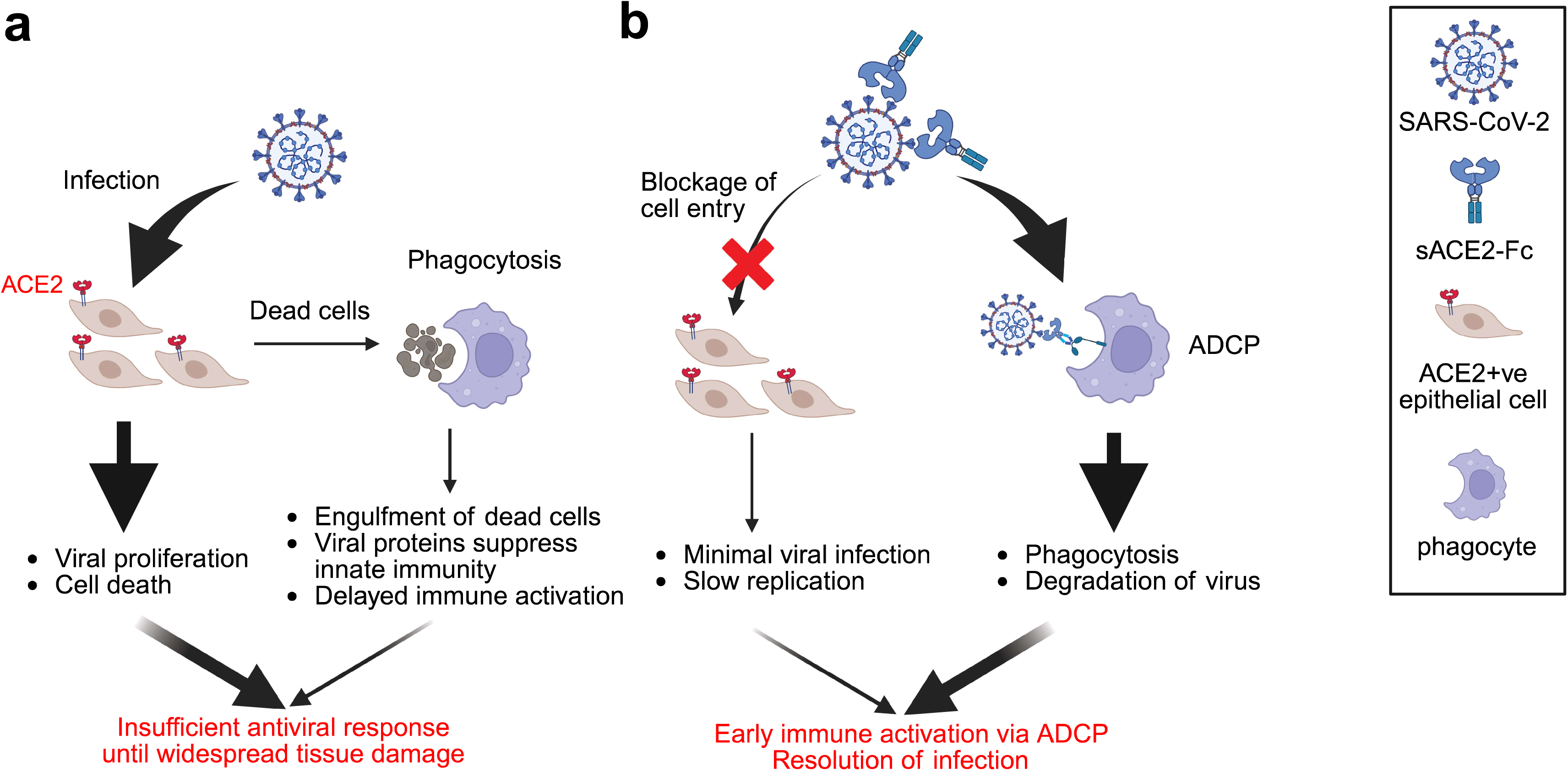
Proposed mechanisms of action of IN sACE2-Fc decoy in preventing SARS-CoV-2 infection. a, **b** Schematics illustrating the actions and outcomes of SARS-CoV-2 infection, in the absence (**a**) and presence (**b**) of IN delivered sACE2-Fc decoys. The figure was generated in Biorender.

In a post-infection context, respiratory viruses would have propagated extensively at the primary infection site when symptoms arise, which makes IN neutralization relatively ineffective to clear the high viral burden and the resultant systematic inflammation [48]. Therefore, the therapeutic activity of either local or systemic neutralization at post-infection stage is bound to be restricted, which is intrinsically inherited in all decoy/antibody therapies against respiratory viral infections. Yet, transfusion of convalescent plasma or infusion of neutralizing mAbs in clinical practice have been shown to reduce the mortality in COVID-19 [49, 50], and IV administration of B5-D3 as therapy was also shown effective in improving the viral burden and survival in our aged mouse experiment (Fig. 2a–e). Therefore, the therapeutic value of our decoy strategy does exist when a “systematic” route of administration (*e.g.*, IV infusion) is chosen, and improvement in disease severity should thus be expected.

Our study has several limitations. First, we didn’t examine the potential of IgM-based sACE2 decamers, which showed higher avidity to spikes and greater potency for viral neutralization in previous studies [46, 51, 52]. It is promising that our B5-D3 design would benefit from switching to the IgM isotype, whereas the distinct biological features imposed by IgM Fc, including short serum half-life and restricted tissue penetration [53], may complicate the analysis design and diverge our study focus. Moreover, although no sign of B5-D3-specific immune responses was observed in our AAV-injected mice, its immunogenicity and potential risk for antibody-dependent enhancement (ADE) should be thoroughly evaluated to further develop the decoy strategy for human use. Furthermore, despite that K18-hACE2 mice are broadly used and demonstrate the best susceptibility to SARS-CoV-2 infection among established hACE2-transgeic mouse models [54], hACE2 expression in these mice shows a distinct pattern that may not reflect human physiology [31, 55]. Further study on hACE2 decoy using animals with physiological level of hACE2 expression and humanized immune system that can support stable engraftment of myeloid lineages would be warranted.

In conclusion, we present a rationally designed sACE2-Fc decoy with minimal mutagenesis (B5-D3) and provide compelling evidence and insights into the immune mechanism supporting its potent prophylactic efficacy. Intranasal prophylactic administration of B5-D3 not only neutralizes SARS-CoV-2 but also redirects the virus toward phagocytic clearance, enabling early immune engagement and complete protection. These findings provide a mechanistic basis for decoy-based antiviral strategies and offer a promising approach to combat current and future airborne viral threats. Further studies may aim to develop approaches to enhance the rapid local immune engagement to restrict early viral propagation. Additionally, regimen refinements are needed to enhance stability and functionality of decoy-based treatments before their clinical translation and extension to a broader range of respiratory pathogens.

## MATERIALS AND METHODS

### Plasmid construction

The coding sequence of human ACE2 was cloned into the pGEM-T easy vector (Promega) and underwent site-directed mutagenesis [10–12, 17]. The sACE2 and human IgG1 hinge-Fc regions (aa 216-447) were assembled via overlapping PCR. These constructs, along with 6xHis-tagged versions, were inserted into the HDM-SARS2-Spike-delta21 vector (Addgene #155130) to generate HDM-CMV-sACE2(-Fc)-his plasmids. L234A/L235A (LALA) in hIgG1 were introduced to generate HDM-CMV-sACE2-Fc-LALA-his plasmids. Various sACE2-Fc fragments were then subcloned into pAAV-nEFCas9 vector (Addgene #87115) to generate AAV-nEF-sACE2-Fc plasmids. SARS-CoV-2 spike variants with or without an HA tag fused to the C-terminal were synthesized and inserted into the HDM vector [56].

### Protein structure visualization

The crystal structure of SARS-CoV-2 spike receptor-binding domain bound with ACE2 (6M0J)was downloaded from the Protein Data Bank (PDB, https://www.rcsb.org/structure/6m0j) [57]. Color-labeling of individual amino acids was performed on the PDB website. For structural overlapping analysis of WT sACE2 and B5-D3 (aa 18-740), protein structures were predicted using online AlphaFold 3 server (https://alphafoldserver.com/) [22]. PyMOL was utilized for root mean square deviation (RMSD) calculations and structural visualization.

### Cell culture

293T, Vero E6, Calu-3, and THP-1 cells were obtained from the American Type Culture Collection and incubated at 37 °C with 5% CO_2_. Specifically, 293T, Vero E6, and Calu-3 cells were maintained in Dulbecco’s Modified Eagle Medium (DMEM, Gibco) supplemented with 10% fetal bovine serum (FBS, Gibco) and 1% Penicillin-Streptomycin (PS, Gibco). THP-1 cells were cultured in Roswell Park Memorial Institute 1640 medium (RPMI, Gibco) with similar supplements. THP-1 cells were differentiated into M0 macrophages using 50 nM phorbol 12-myristate 13-acetate (PMA) for 48 h, followed by a 24 h rest. For M1 macrophage differentiation, post-PMA treatment cells were stimulated with 10 ng/mL lipopolysaccharide and 20 ng/mL interferon (IFN)-γ for 24 h. Expi293 cells (Gibco) were cultured following the manufacturer’s instructions.

### Immunofluorescence staining of hACE2-293T and hACE2-Calu-3

Cells were fixed, permeabilized, and blocked with 10% Normal Goat Serum (Invitrogen). ACE2 was stained with a primary antibody (Abcam #ab15348) followed by an Alexa Fluor 594-conjugated secondary antibody (Invitrogen #A-21442). Cells were counterstained with Hoechst 33342 (Thermo Scientific) and examined under Nikon Ti2-E Inverted Fluorescence Microscope.

### Lentivirus packaging and transduction

293T cells were seeded at 80% confluence and transfected with psPAX2 (Addgene #12260), pMD2.G (Addgene #12259), and transfer plasmid pWPI-IRES-Puro-Ak-ACE2-TMPRSS2 (Addgene #154987) using polyethylenimine (PEI). Lentivirus-containing medium was harvested 72 h post-transfection, filtered through a 0.45 µM filter, concentrated, and stored at −80°C. For transduction, 293T or Calu-3 cells were exposed to the concentrated lentivirus with 8 μg/mL polybrene for 24 h to obtain human ACE2-overexpressing cell lines (hACE2-293T and hACE2-Calu-3 respectively).

### Pseudovirus packaging, titration, and infection

Pseudoviruses were packaged in 293T cells using pCDH-EF1a-eFFly-eGFP (Addgene #104834) and spike-encoding plasmids, following a similar protocol to that of lentivirus. Post-packaging, pseudoviral particles were titrated using the Lenti-X qRT-PCR Titration Kit (Takara #631235) and used to infect target cells in the presence of 8 µg/mL polybrene. Infectivity was assessed via a luciferase assay (Promega #E1501).

### Protein production and purification

293T and Expi293 cells were transfected with HDM-CMV-sACE2(-Fc)(-LALA)-his plasmids using PEI and ExpiFectamine 293 Transfection Kit (Gibco) respectively. Culture supernatants were collected after 72 h and 5 days post-transfection respectively. The 293T supernatant was assessed for ACE2 and IgG1 levels using enzyme-linked immunosorbent assay (ELISA) kits (Abcam #ab235649; Invitrogen #BMS2092), and ACE2 activity was measured with a fluorometric assay (Abcam # ab273297). The Expi293 supernatant underwent Ni-NTA Agarose purification, followed by elution and buffer exchange to phosphate-buffered saline (PBS, pH 7.4). Protein concentration and integrity were verified using the Bradford method, ELISA, and sodium dodecyl sulfate-polyacrylamide gel electrophoresis (SDS-PAGE).

### *In vitro* pseudovirus neutralization assay

Conditioned media containing sACE2, sACE2-Fc or sACE2-Fc-LALA proteins were diluted serially, mixed with pseudovirus (4 × 10^9^ copies), and incubated at room temperature for 30 minutes [19]. The mixture was added to hACE2-293T cells in 96-well plates with duplicates with 8 µg/mL polybrene. Transduction efficiency was assessed 48 h later via green fluorescent protein (GFP) imaging and/or luciferase assays.

### Reporter-based *in vitro* ADCC and ADCP assays

The *in vitro* antibody-dependent cell-mediated cytotoxicity (ADCC) and antibody-dependent cellular phagocytosis (ADCP) activities of B5-D3(-LALA) were measured using Jurkat-Lucia NFAT-CD16 and Jurkat-Lucia NFAT-CD32 cells (InvivoGen) respectively according to the manufacturer’s instructions. 293T cells transfected with pBOB-CAG-SARS-CoV-2-Spike-HA (Addgene #141347) acted as target cells. Target cells were co-incubated with reporter cells and serially diluted B5-D3(-LALA) at 37°C for 1 h. Luciferase expression indicating CD16 and CD32 signaling was measured using QUANTI-Luc (InvivoGen).

### ADCP of pseudovirus in THP-1 and THP-1-derived macrophages

SARS-CoV-2 pseudovirus (Wuhan-Hu-1, 8 × 10^8^ copies in 10 µL) was mixed with 50 µL of B5-D3 or control proteins (20 µg/mL) and added to THP-1, M0, or M1 cells cultured in a µSlide 18 Well iBITreat chamber slide. Here THP-1 cells were attached on the collagen-coated iBITreat chamber slide. After 1, 3, 6, or 18 hours of incubation for phagocytosis, cells were fixed with ice-cold methanol for 10 minutes, blocked with 10% normal goat serum for 1 hour at room temperature, and immunostained sequentially for human IgG-Fc (Abcam #ab98596), HIV-1 p24 (Invitrogen #PA5-81773), and lysosomal associated membrane protein 1 (LAMP1) (Abcam #ab25630). Secondary antibodies were applied (Invitrogen #A-21200 and #A-31573), and nuclei were stained with Hoechst. Imaging was conducted using a Leica SP8 confocal microscope. p24 immunofluorescence intensity and colocalization with LAMP1 (Manders’ coefficient) were analyzed using ImageJ and the JACoP plugin.

For RNA extraction, M0 cells were washed once with PBS and added with TRIzol (Invitrogen) after 6 h incubation with pseudovirus with/without B5-D3 (*n* = 3 in triplicate wells).

### Western blot for spike cleavage detection

SARS-CoV-2 spike-HA tagged pseudovirus (4 × 10^9^ copies) was incubated with M0/M1 macrophages or hACE2-293T cells for 6 h, with or without sACE2-Fc proteins. After incubation, cell lysates were processed through SDS-PAGE and transferred to polyvinylidene difluoride membranes. The membranes were blocked, incubated overnight with anti-HA (Merck Millipore #05-904) and anti-β-actin (Santa Cruz #sc-47778) primary antibodies, then with horseradish peroxidase (HRP)-conjugated secondary antibodies (Cell Signaling Technology #7076). Signals were detected using the Amersham ECL select kit on a Bio-Rad ChemiDoc MP system.

### AAV vector packaging and purification

As described previously [58], AAV vectors were produced in 293T cells transfected with AAV-nEF-sACE2-Fc, pAdDeltaF6 (Addgene #112867), and pAAV2/8 (Addgene #112864) plasmids using PEI. AAV particles were harvested from supernatants with Polyethylene Glycol (PEG) 8000 and from cell lysates via freeze-thaw cycles and benzonase digestion, then purified using Iodixanol density gradient ultracentrifugation. AAV particles were concentrated and measured using a qPCR AAV Titer Kit (Applied Biological Materials #G931).

### SARS-CoV-2 virus

Experiments with live SARS-CoV-2 were performed at the BSL-3 core facility (LKS Faculty of Medicine, HKU). The BetaCoV/Hong Kong/VM20001061/2020 virus, here regarded as the wild type strain of SARS-CoV-2 (Wuhan-Hu-1), was isolated from the nasopharyngeal aspirate and throat swab of a confirmed patient with COVID-19 in Hong Kong (GISAID identifier EPI_ISL_412028). The SARS-CoV-2 variants were isolated from clinical specimens in Hong Kong. Stock viruses were prepared with Vero E6 cells cultured in infection medium (DMEM supplemented with 2% FBS and 1% PS).

### Median tissue culture infectious dose (TCID_50_) assay

Vero E6 cells pre-seeded in 96-well plates were infected with serially diluted virus stocks or mouse lung homogenates in infection medium. After 72 h incubation, cytopathic effects (CPEs) were observed under a microscope to calculate titers using the Reed–Muench method.

### Plaque-reduction neutralization test (PRNT) assay

Casirivimab (#C100P) and hIgG1 isotype (clone 4F17, #PA007125) were purchased from Syd Labs, USA) and used as controls. Purified B5-D3 and control proteins were serially diluted and incubated with 50 PFU of SARS-CoV-2 (Wuhan-Hu-1, Delta, Omicron BA.5, BQ.1.22, and XBB.1.5) for 1 h at room temperature, followed by addition to Vero E6 cells seeded in 6-well plates. After incubation, cells were overlaid with agarose, fixed with formalin, and stained with crystal violet. Plaque counts were used to calculate percentage neutralization and half maximal inhibitory concentration (IC_50_) values. The experiments were carried out in duplicates.

### Animal experiments

The K18-hACE2 mice (B6.Cg-Tg(K18-ACE2)2Prlmn/J) [31] were purchased from the Jackson Laboratory (Bar Harbor, ME, USA). Experiments on AAV or protein-only administration in mice were carried out in the Animal Holding Core of the School of Biomedical Sciences, CUHK. Experiments involving SARS-CoV-2 infection in K18-hACE2 mice were conducted within the confines of the Biosafety Level 3 (BSL-3) core facility located at the Li Ka Shing Faculty of Medicine, HKU. Experiments were conducted according to ethical practices to minimize animal distress.

### AAV-mediated sACE2-Fc overexpression in mice

Male K18-hACE2 mice [31] at the age of 2 months were injected intravenously with 1 × 10^11^ GC of AAV-nEF-sACE2-Fc. Blood samples were collected periodically for analysis. Following euthanasia, tissues were harvested for further DNA and histological analysis. sACE2-Fc concentrations in sera and RAS metabolites were quantified using specific ELISA kits for Human IgG1, Mouse Renin 1 (Invitrogen #EMREN1), Angiotensin II (LifeSpan BioSciences #LS-F523), and Angiotensin 1-7 (LifeSpan BioSciences #LS-F40645).

### SARS-CoV-2 infection in mice

Female K18-hACE2 mice, aged 10-12 months or 2-3 months, were intranasally inoculated with 1×10^4^ plaque-forming unit (PFU) of SARS-CoV-2 Wuhan-Hu-1. Treatment with B5-D3 protein was administered intranasally at 2.5 mg/kg or intravenously at 15 mg/kg, at various time points relative to the viral challenge (6 h before, 24 h before, or 24 h after). Vehicle control groups received PBS 6 h before viral challenge. Survival and weight were monitored daily for 14 days. For the older mice, lung samples were collected from one mouse from each group at 4 days post-infection (dpi) for analysis; younger mice had plasma collected for neutralizing antibody analysis at 14 dpi. Another batch of young mice also received B5-D3(-LALA) protein pre-inoculation, with lungs analyzed post-inoculation for RNA, viral load, and histopathology. A control group of non-infected mice was used to assess baseline effects of B5-D3 on lung tissue.

### Neutralization assay for antibody titration

Vero E6 cells were pre-seeded on 96-well plates 24 h before infection. On the day of infection, the growth medium of the cells was changed to infection medium. The plasma samples were serially 2-fold diluted with infection medium from a starting dilution of 1:10. The plasma was then pre-incubated with 100 TCID_50_ of SARS-CoV-2 for 1 h at room temperature before being inoculated to the seeded Vero E6 cells in quadruplicates. At 72 h after inoculation, CPEs of the cells were observed with optical microscopy. Neutralizing antibody titers against SARS-CoV-2 were expressed as the reciprocal of the highest dilution of plasma showing no CPEs in all 4 wells. Uninfected cell monolayers were used as toxicity control.

### Histology

Mouse tissues were fixed in 10% formalin, embedded in paraffin, and sectioned at 5 μm. Sections of different organs were deparaffinized and underwent hematoxylin and eosin (H&E) staining. For immunohistochemistry (IHC) staining, lung sections underwent antigen retrieval, endogenous peroxidase blocking, and were incubated with primary antibodies against the SARS-CoV/SARS-CoV-2 nucleocapsid protein (Sino Biological #40143-T62) overnight. After washing, sections were stained with the anti-rabbit VECTASTAIN Elite ABC-HRP Kit (Vector Laboratories), developed with 3,3′-Diaminobenzidine (Sigma #D4293), and counterstained with Mayer’s hematoxylin. Stained sections were scanned using Zeiss Axioscan 7 and analyzed using ZEN (blue edition). For measurement of alveolar septal thickness, 10 fields from each lung H&E section and 10 septa from each field were chosen for measurement using ImageJ.

### Quantitative PCR

Quantitative PCR (qPCR) was used to analyze liver genomic DNA and lung RNA from mice. DNA was extracted and qPCR was performed with the TB Green Premix Ex Taq II kit (Takara), normalized to mouse *Gapdh* using the 2^-ΔCt method. RNA was extracted using TRIzol, reverse-transcribed (Applied Biosystems #4368813), subjected to qPCR using the same kit and normalized to mouse *Gapdh* levels. Specific primers are listed in Supplementary Table 2.

### RNA-Seq and data analysis

Total RNA was extracted from mouse lung tissues or THP-1-derived M0 macrophages using TRIzol and processed into transcriptome libraries with the TruSeq RNA Library Prep Kit (illumina). Sequencing was performed on the NovaSeq 6000 or NovaSeq X Plus sequencers (illumina) using a 150-base pair paired-end configuration. Sequencing data were processed with fastp for quality control [59], then aligned to both the mouse (Ensembl GRCm39) and SARS-CoV-2 (NCBI NC_045512v2) genomes using STAR [60] or to the human genome (Ensembl GRCh38) using HISAT2 [61]. Pearson correlation and groupwise comparisons were conducted in R: gene expression was quantified and analyzed for differential expression using DESeq2 [62]; up/downregulated gene enrichment and Gene Set Enrichment Analysis (GSEA) [63] was performed using the clusterProfiler package [64].

### Tracking B5-D3 bio-distribution in mice

B5-D3 were conjugated with Alexa Fluor 750 dye (B5-D3-AF750) as described [46]. In brief, 2 mg/mL solution of B5-D3 protein in 0.15 M NaHCO_3_ was reacted with Alexa Fluor 750 succinimidyl ester (Thermo Fisher Scientific) at room temperature for 1 h. Unreacted dye was removed by dialysis in PBS. All procedures were performed under dimmed light. Female K18-hACE2 mice, aged 2-3 month, were administered intranasally with B5-D3-AF750 (2.5 mg/kg). The mice were imaged at predetermined time points after administration (fluorescence ex = 745 nm, em = 800 nm, auto-exposure setting) using an IVIS Spectrum CT Imager (Perkin Elmer). At the time of euthanasia, 50 μL of urine and blood, the brain, nasal cavity, trachea, lung, heart, liver, spleen, kidney, and urinary bladder samples were excised and imaged. Regions of interest (ROIs) were drawn, and average radiance (p/s/cm²/sr) was measured. All images were processed using Living Image software (Perkin Elmer) and the same fluorescence threshold was applied for group comparison.

### Flow cytometry analysis and confocal microscopic imaging of BALF cells

Mice were sacrificed via anesthetics overdose. Bronchoalveolar lavage was performed by intratracheally rinsing the lungs with 1 mL of ice-cold Hanks’ Balanced Salt Solution (HBSS, Gibco) containing 100 μM ethylenediaminetetraacetic acid for four repeats. Bronchoalveolar lavage fluid (BALF) was then centrifuged and treated with ammonium-chloride-potassium red blood cell lysing buffer. For multicolor flow cytometry, cell pellets were washed with PBS and stained with the Fixable Viability Stain 440UV dye (BD #566332). Next, the cells were blocked with CD16/CD32 monoclonal antibody (Invitrogen #14-0161-85) and stained with antibodies targeting the following molecules: CD45 (BD #568336), Siglec-F (BD #564514), CD11b (BD #612800), CD11c (BD #751265), Ly6G (BD #563005), I-A/I-E major histocompatibility complex class II (MHC-II) (BD #750171), F4/80 (BD #570288), Ly6C (BD #755198), and CD3 (BD #555275). Stained BALF cells were analyzed using the BD FACSymphony A5.2 SORP Flow Cell Analyzer, and the results were analyzed using FlowJo v10.10.

For microscopic inspections, BALF cells were collected from mice, seeded in poly-d-lysine-coated chamber slides and stained with antibodies targeting Siglec-F (BD #564514) and human IgG-Fc (Abcam #ab98596). Secondary antibody (Invitrogen #A-11006) was applied. Stained BALF cells were then counterstained with Hoechst and examined by confocal microscopy (Leica TCS SP8).

### Statistical Analysis

Assays including *in vitro* neutralization, PRNT, ADCC, and ADCP were conducted in technical duplicates. Results were analyzed in GraphPad Prism version 9 using nonlinear regression to calculate IC_50_ or half maximal effective concentration (EC_50_) values. Transcriptomic analyses were performed using R, with details provided in figure captions. All other statistical analyses utilized GraphPad Prism version 9 with a significance threshold set at *p* value (*p*) < 0.05.

## DECLARATIONS

### Ethics approval and consent to participate

All animal procedures were ethically approved by The Chinese University of Hong Kong (CUHK)’s Animal Experimentation Ethics Committee (approval number: 20-226-MIS) and The University of Hong Kong (HKU)’s Committee on the Use of Live Animals in Teaching and Research (approval number: 5511-20).

### Availability of data and materials

All data associated with this study are available in the main text or the supplementary materials. The RNA-seq data generated in this study have been deposited in the NCBI Sequence Read Archive database under accession code PRJNA1054508. Constructs of diverse sACE2-Fc mutants and SARS-CoV-2 spikes are available upon request after completion and approval of a material transfer agreement by contacting.

### Authors’ contributions

Jingyi W. and J.L. constructed the sACE2-Fc mutants and performed characterization analysis; A.W.C. performed the PRNTs and data analysis; Jingyi W., A.W.C., and J.L. performed the mouse infection experiments and data analysis; B.L., J.Q. and J.R. produced recombinant sACE2-Fc proteins; Jingyi W. and Junkang W. performed RNA-Seq analysis; Jingyi W. and J.Q. performed protein labeling, *in vivo* tracing of labeled protein, and flow cytometry analysis of BALF cells; J.L. performed THP-1 and Calu-3 experiments and confocal microscopic analysis. Junkang W. and J.L. performed protein structure prediction and visualization. Jingyi W., J.L., L.L.P. and B.F. conceived the project, designed experiments, and wrote the manuscript. Y.X., T.B., L.L.P. and B.F. revised the manuscript. All authors read and approved the final manuscript.

## Acknowledgements

We thank the Chinese University of Hong Kong (CUHK) and the University of Hong Kong (HKU) research platforms for assistance in animal experimentation (the Laboratory Animal Service Center at CUHK and the Centre for Comparative Medicine Research at HKU) and histological analysis (Department of Pathology, HKU and Core Laboratory in the School of Biomedical Sciences, CUHK).

## Supplementary Information for

**Supplementary Fig. 1.**
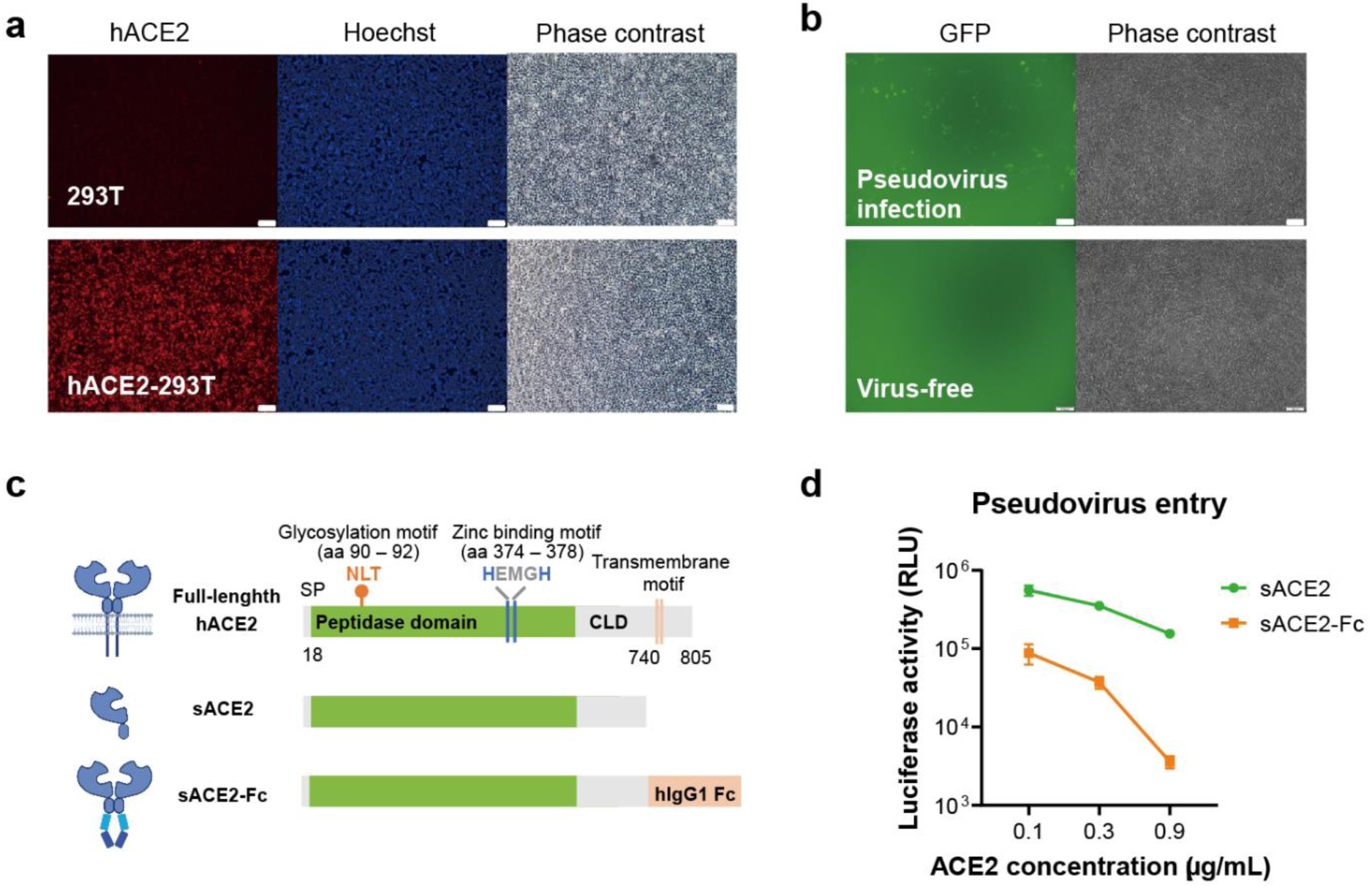
Establishment of the pseudoviral infection platform and generation of sACE2 decoys. **a** Microscopic images show the expression of full length human ACE2 (hACE2) in stable hACE2-293T cells established by lentiviral transduction (Scale bar = 100 μm). Immunostaining was performed using antibody specific to hACE2 (Abcam # ab15348). **b** Representative fluorescence and phase contrast images showing GFP expression in hACE2-293T cells with and without infection by pseudovirus carrying the Wuhan-Hu-1 spike protein (Scale bar = 100 μm). **c** Schematic diagrams of hACE2 (top), sACE2 (middle), and sACE2-Fc (bottom) molecules indicating important amino acid positions (90-92 and 374-378) for glycosylation and zinc binding. aa, amino acid. **d** Line chart comparing the neutralization efficiencies of sACE2 (green) and sACE2-Fc (orange) against Wuhan-Hu-1 pseudovirus expressing luciferase, measured in relative luminescence units (RLU). Data are presented as mean ± SD from duplicate experiments.

**Supplementary Fig. 2.**
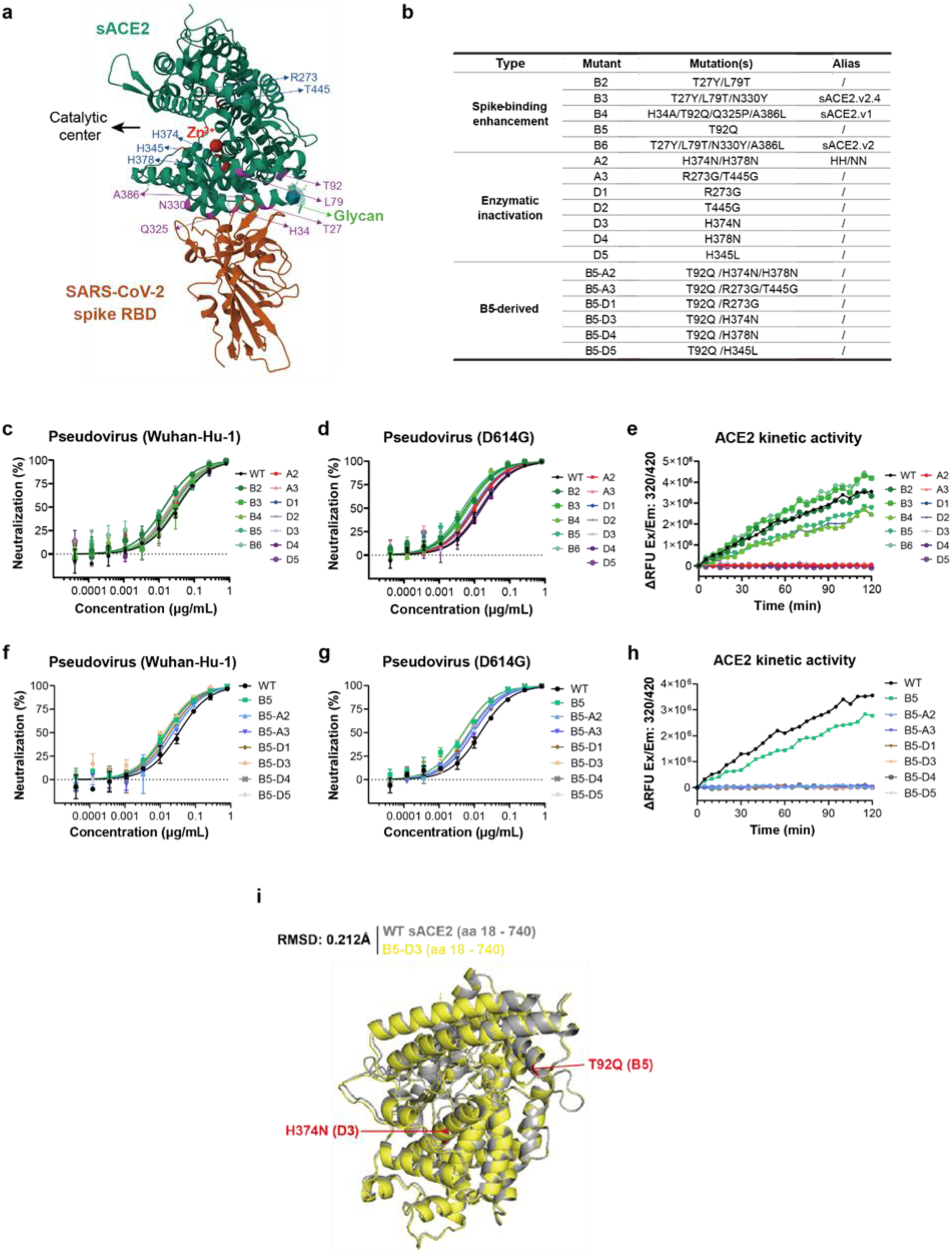
Engineering and characterization of enhanced sACE2 decoys. **a** 3D structural model of ACE2 (green) complexed with the SARS-CoV-2 spike receptor-binding domain (RBD, brown), highlighting mutations for spike-binding enhancement (magenta) and enzymatic inactivation (blue). Structures were adapted from Protein Data Bank (PDB ID: 6M0J). **b** List of mutations in sACE2 sequences tested for enhanced binding or enzymatic inactivation. **c**, **d** Neutralization assay results for WT sACE2-Fc and mutants (B2 to B6, A2, A3, and D1 to D5) against Wuhan-Hu-1 (**c**) and D614G (**d**) pseudoviruses. **e** Kinetic curves showing the ACE2 enzymatic activities of WT sACE2-Fc and B2 to B6, A2, A3, D1 to D5 mutants. **f**, **g** Neutralization results for WT sACE2-Fc and mutants (B5, B5-A2, B5-A3, B5-D1, B5-D3, B5-D4 and B5-D5) against Wuhan-Hu-1 (**f**) and D614G (**g**) pseudoviruses. **h** Kinetic curves showing the ACE2 enzymatic activities of WT sACE2-Fc and B5, B5-A2, B5-A3, B5-D1, B5-D3, B5-D4 and B5-D5 mutants. **i** Conformational comparison between WT and B5-D3 sACE2. The 3D structures of WT and B5-D3 sACE2 (aa 18–740) were predicted using AlphaFold 3 and superimposed for direct comparison. The WT sACE2 structure is coloured in grey, while the B5-D3 variant is highlighted in yellow. Mutations in the B5-D3 structure are specifically marked in red. Data are presented as mean ± SD from duplicate experiments.

**Supplementary Fig. 3.**
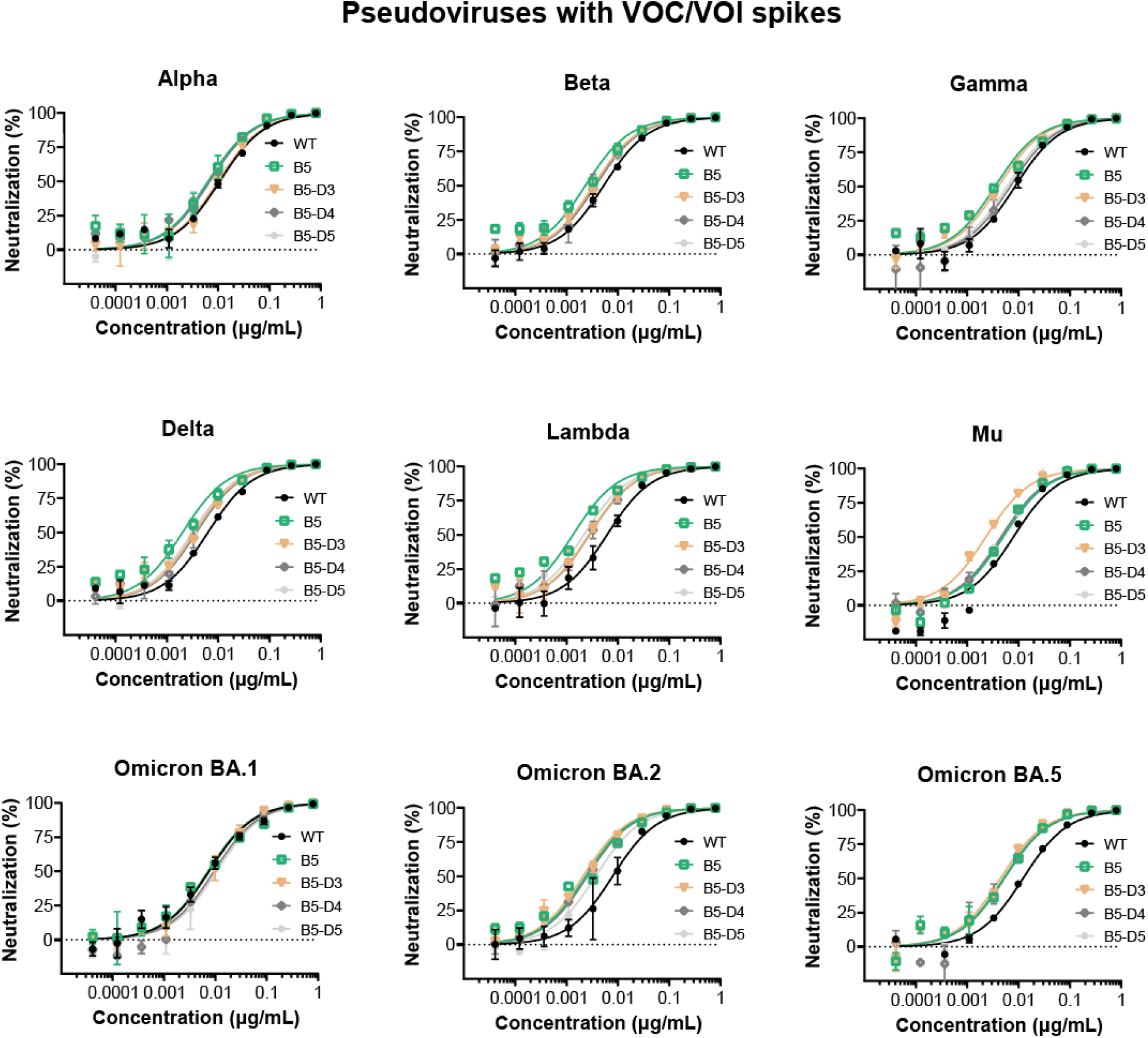
Broad-spectrum neutralization against pseudoviruses of SARS-CoV-2 VOC/VOI strains by sACE2-Fc candidate mutants. Graphical representation of dose-response curves in neutralization assays against pseudoviruses bearing spikes from various SARS-CoV-2 VOCs and VOIs. Data are presented as mean ± SD from duplicate experiments.

**Supplementary Fig. 4.**
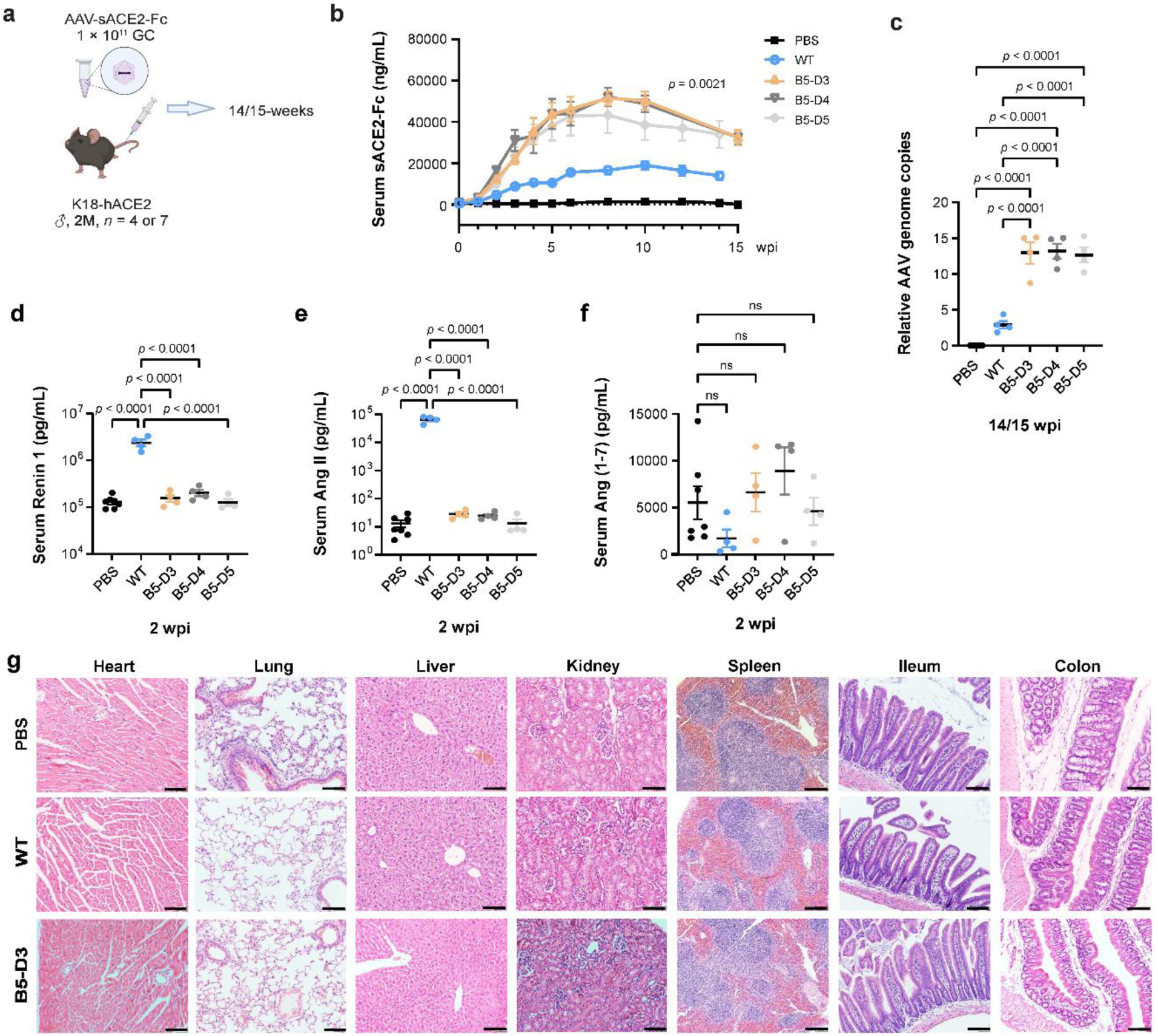
AAV-delivered prolonged overexpression of WT sACE2-Fc and candidate mutants in K18-hACE2 mice. **a** Schematic of AAV administration. Male K18-hACE2 mice aged 2 months received tail vein injections of either PBS (black; *n* = 7) or AAV carrying WT sACE2-Fc (blue), B5-D3 (orange), B5-D4 (dark grey), or B5-D5 (light grey) (*n* = 4 each) at a dose of 1 × 10^11^ GC. Mice were observed for up to 15 weeks and then sacrificed for tissue analysis. wpi, weeks post-injection; M, month. **b** Serum concentrations of sACE2-Fc were quantified via ELISA using antibodies specific to hIgG1. **c** Quantification of AAV genomes in the mouse livers at the observation endpoint. Results shown are from qPCR of genome DNA normalized to mouse *Gapdh.* **d**–**f** Concentrations of renin (**d**), Ang II (**e**), and Ang (1-7) (**f**) in sera at 2 wpi, measured via metabolite-specific ELISA. ns, not significant with *p* value (*p*) ≥ 0.05. **g** Representative H&E staining of heart, lung, liver, kidney, spleen, ileum, and colon tissues from mice in the PBS, WT sACE2-Fc, and B5-D3 treatment groups (scale bar = 50 μm). Data are presented as mean ± SEM. Statistical analyses were performed using one-way analysis of variance (ANOVA) and Tukey’s multiple comparisons test following ANOVA.

**Supplementary Fig. 5.**
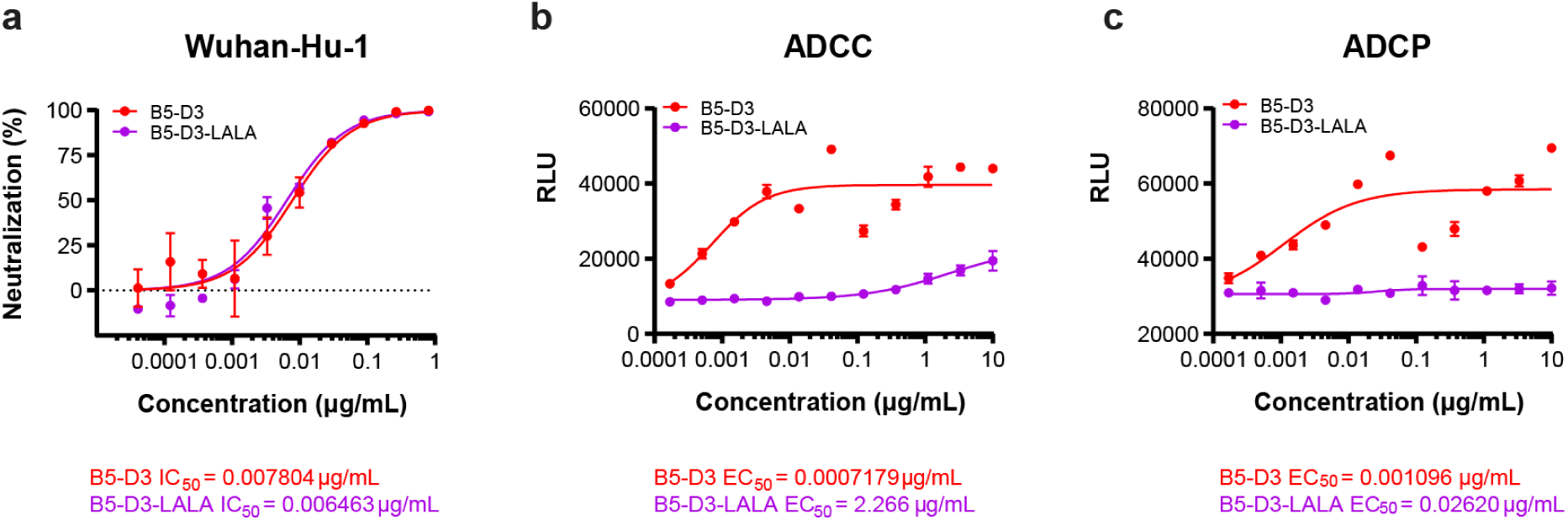
Comparison between B5-D3 and B5-D3-LALA in *in vitro* neutralization against the SARS-CoV2 pseudovirus and Fc-mediated effector functions. **a** Neutralization capability of B5-D3-LALA was compared to B5-D3 using an *in vitro* pseudovirus neutralization assay. IC_50_ values are indicated for each version. **b**, **c** Fc-mediated effector functions of B5-D3 and B5-D3-LALA were assessed using luciferase-based reporter cell lines. Dose-response curves and half maximal effective concentration (EC_50_) values for antibody-dependent cellular cytotoxicity (ADCC) (**b**) and antibody-dependent cellular phagocytosis (ADCP) (**c**) are displayed. Data are presented as mean ± SD from duplicate experiments.

**Supplementary Fig. 6.**
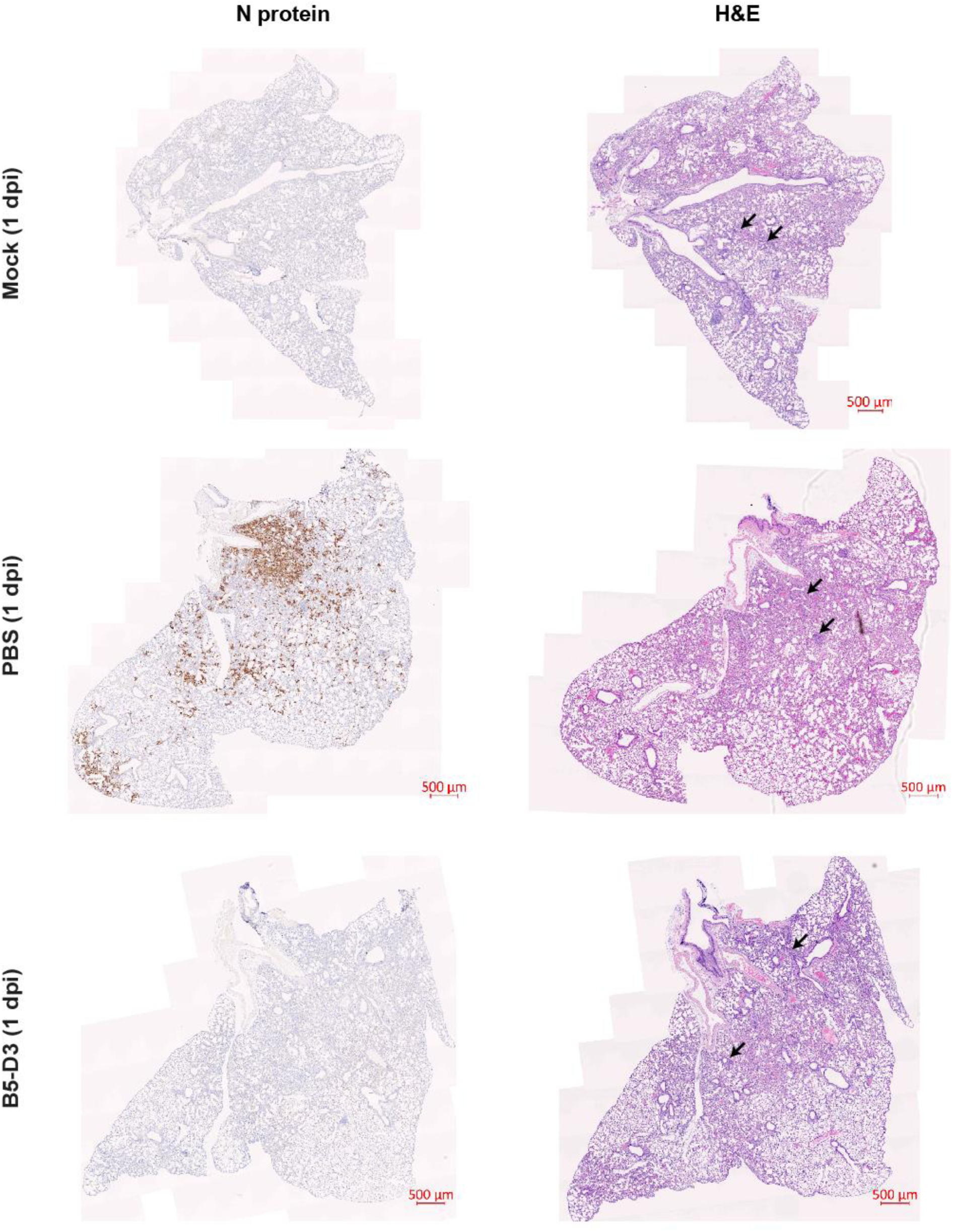
Entire sections for histological examination at 1 dpi. Fixed lung tissues were sectioned and stained for histological examination. IHC staining for viral N protein (left panels, SinoBiological # 40143-T62) and H&E staining for tissue damage (right panels) are displayed. Black arrows in H&E photos indicate areas of alveolar thickening. Scale bar = 500 μm.

**Supplementary Fig. 7.**
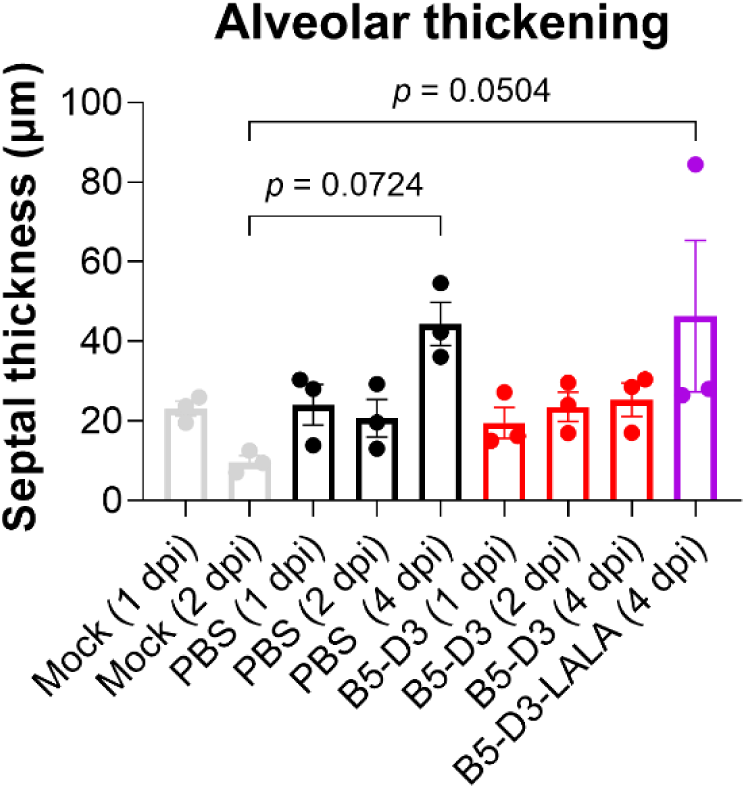
Thickening of alveolar septum in K18-hACE2 mice after SARS-CoV-2 challenge. Lung septum thickness of K18-hACE2 mice in Fig. 3 was measured from H&E staining. Data are presented as mean ± SEM, and statistical significance was determined by Tukey’s multiple comparisons test.

**Supplementary Fig. 8.**
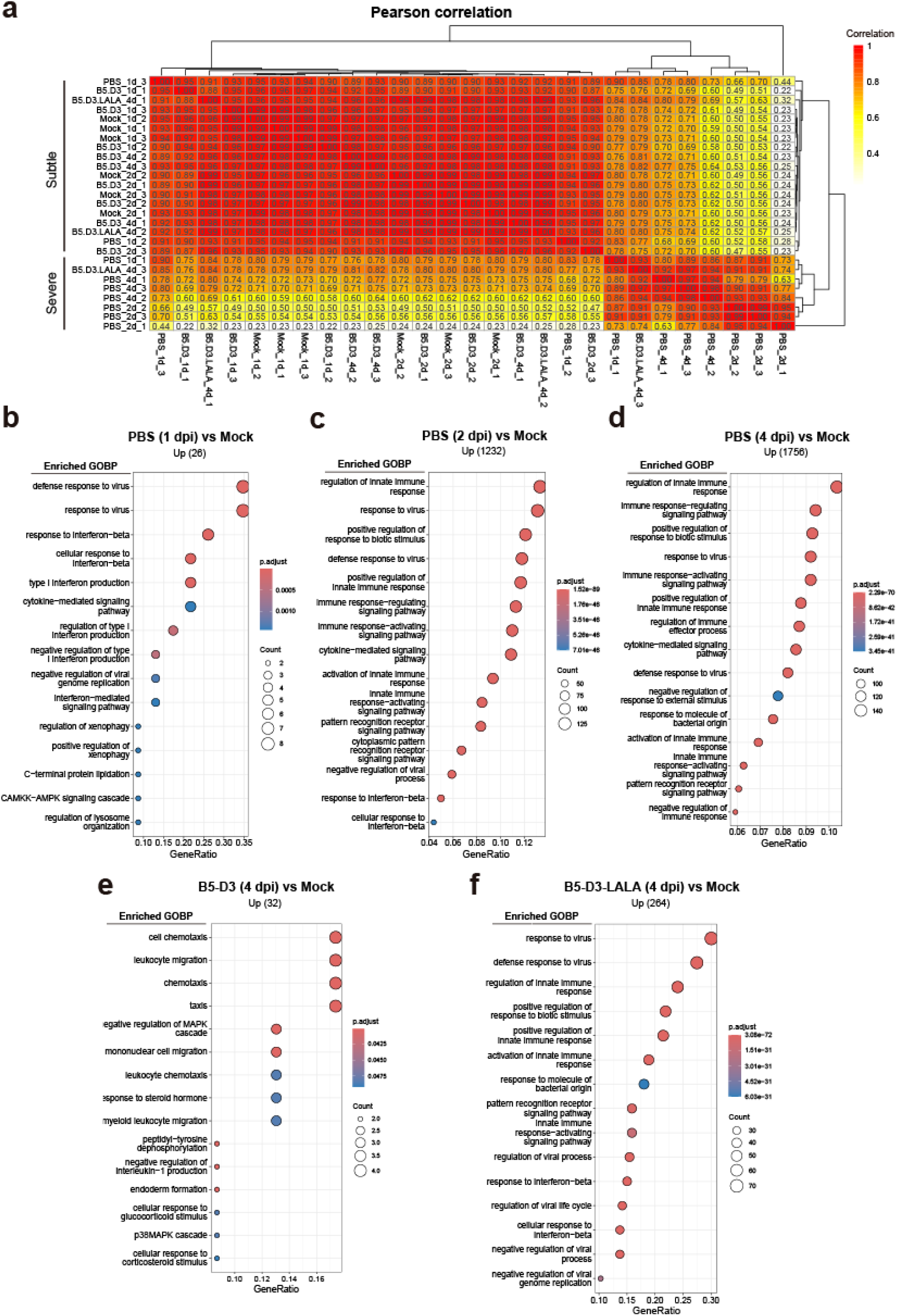
RNA-Seq analysis of K18-hACE2 mouse lungs with different pretreatments upon SARS-CoV-2 challenge. **a** Bulk RNA-seq was conducted on lung homogenates from K18-hACE2 mice under mock treatment (*n* = 6) or post-viral challenge at 1, 2 and 4 dpi with each time point (*n* = 3). Pearson correlation analysis was executed on 27 samples using counts per million (CPM) data, with each cell displaying the Pearson correlation coefficient color-coded for visual ease. **b**–**f** GOBP enrichment analysis identifies biological processes enriched in up-regulated genes from comparisons at 1 (**b**), 2 (**c**), and 4 dpi (**d**) for PBS, 4 dpi for B5-D3 (**e**), and 4 dpi for B5-D3-LALA (**f**) versus the mock control, with top 15 significant terms displayed. *p*.adjust, adjusted *p* value; CAMKK, calmodulin-dependent protein kinase; AMPK, 5’ adenosine monophosphate-activated protein kinase. Benjamin–Hochberg method was used for FDR adjustment.

**Supplementary Fig. 9.**
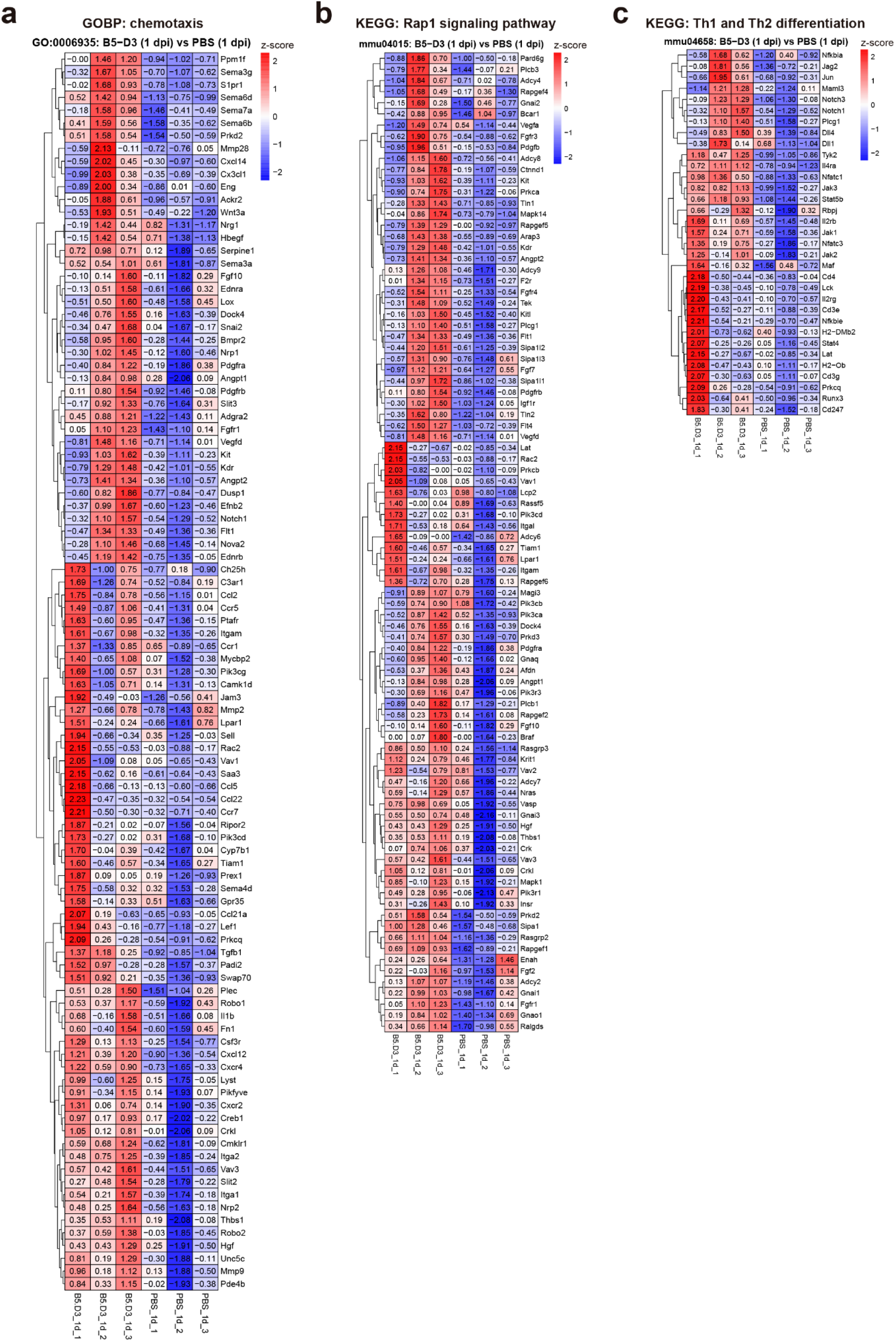
Leading-edge subsets in GSEA. **a**–**c** Z-score plots show the relative level of gene expression in the leading edge subsets from GSEA comparing B5-D3 vs PBS at 1 dpi, corresponding to chemotaxis (**a**), Rap1 signaling pathway (**b**), and Th1 and Th2 differentiation (**c**).

**Supplementary Fig. 10.**
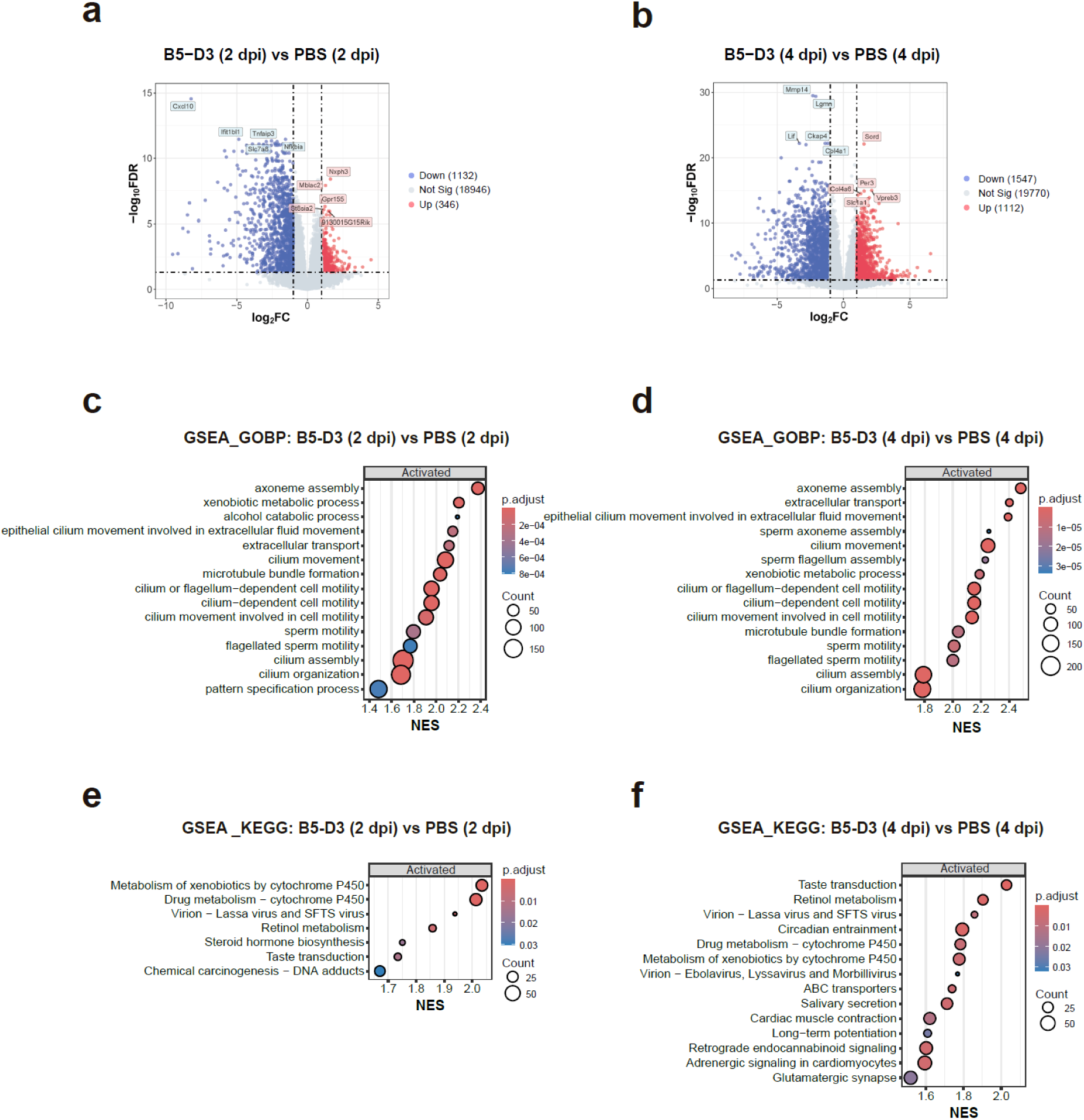
Transcriptomic comparisons between B5-D3 and PBS pretreatments in K18-hACE2 mice upon SARS-CoV-2 challenge. a, **b** DGE analysis between B5-D3 and PBS groups at 2 (**a**), and 4 dpi (**b**) showing up-regulated and down-regulated genes visualized in volcano plots. **c**, **d** GSEA of GOBPs significantly activated in B5-D3 groups compared to PBS groups at 2 dpi (**c**) and 4 dpi (**d**), with top 15 most significant terms displayed. **e, f** GSEA of KEGG pathways significantly activated in B5-D3 groups compared to PBS groups at 2 dpi (**e**) and 4 dpi (**f**), with all significant terms (**e**) and top 15 most significant terms (**f**) displayed. SFTS, Severe Fever with Thrombocytopenia Syndrome. Benjamin–Hochberg method was used for FDR adjustment.

**Supplementary Fig. 11.**
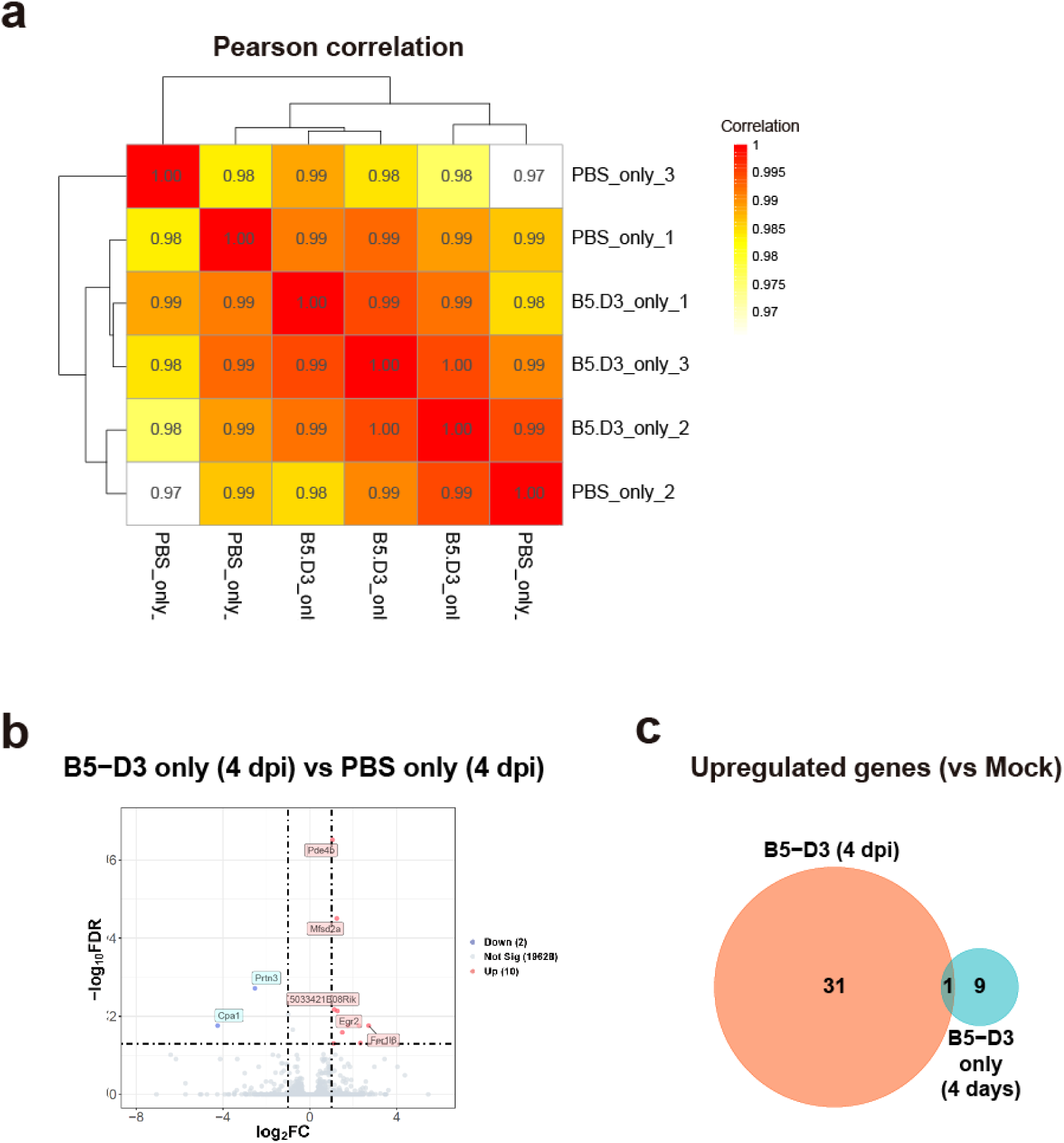
Minimal transcriptomic alterations in lungs after IN B5-D3 administration without viral challenge. **a** Pearson correlation analysis for lung tissues collected 4 days post-administration of IN B5-D3 or PBS in female K18-hACE2 mice (*n* = 3), depicting correlation coefficients. **b** Volcano plot showing differentially expressed genes between the B5-D3 treated and control groups, with significant changes marked. **c** Venn diagram showing overlaps among the upregulated genes in **b** and in B5-D3 (4 dpi) in Fig. 4**c**.

**Supplementary Fig. 12.**
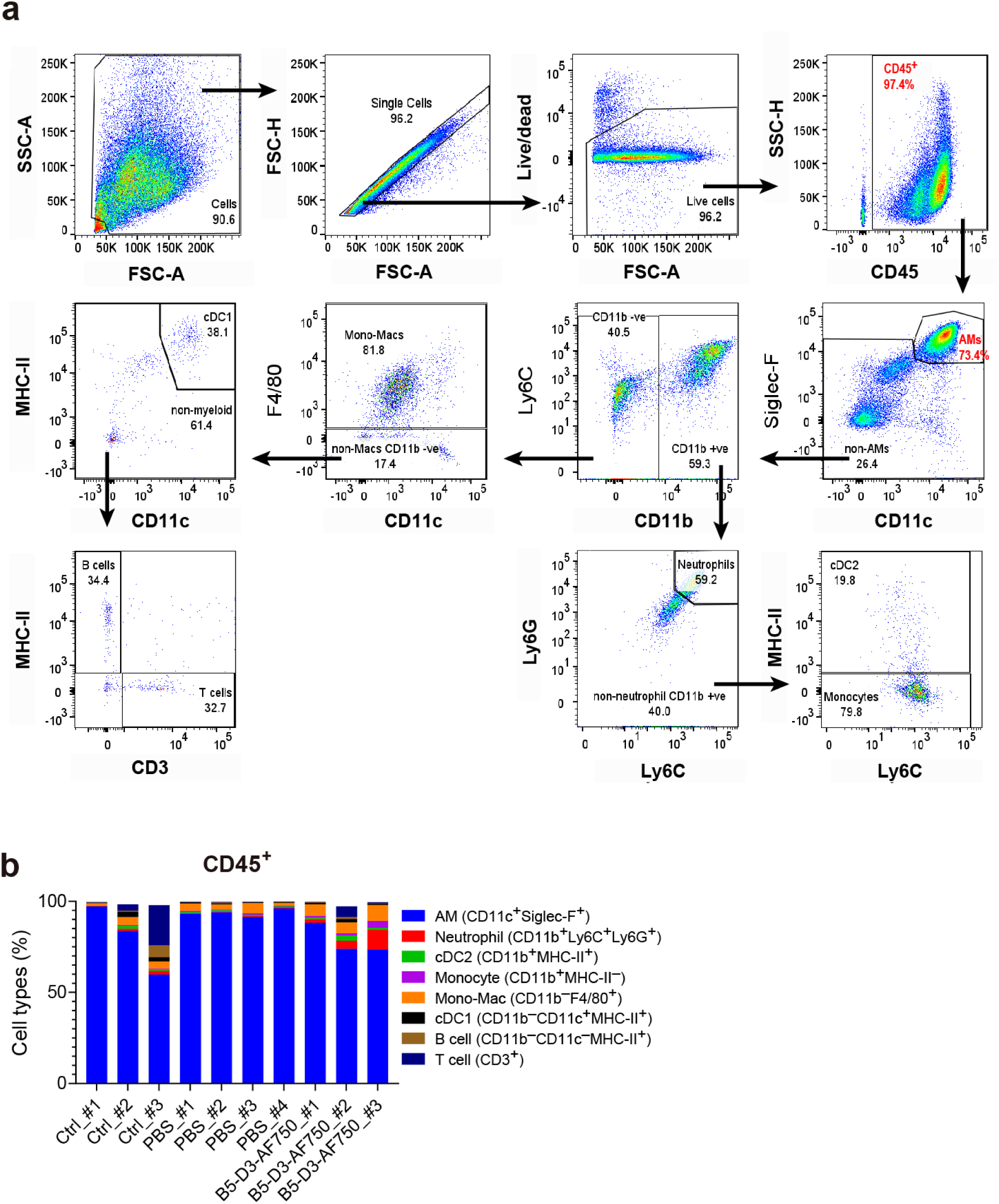
Flow cytometry analysis of mouse BALF cells. **a** Flow cytometric gating strategy for BALF cells. **b** Percentages of individual cell types in CD45^+^ BALF cells collected from individual animals. Ctrl group received no treatment before sacrifice.

**Supplementary Fig. 13.**
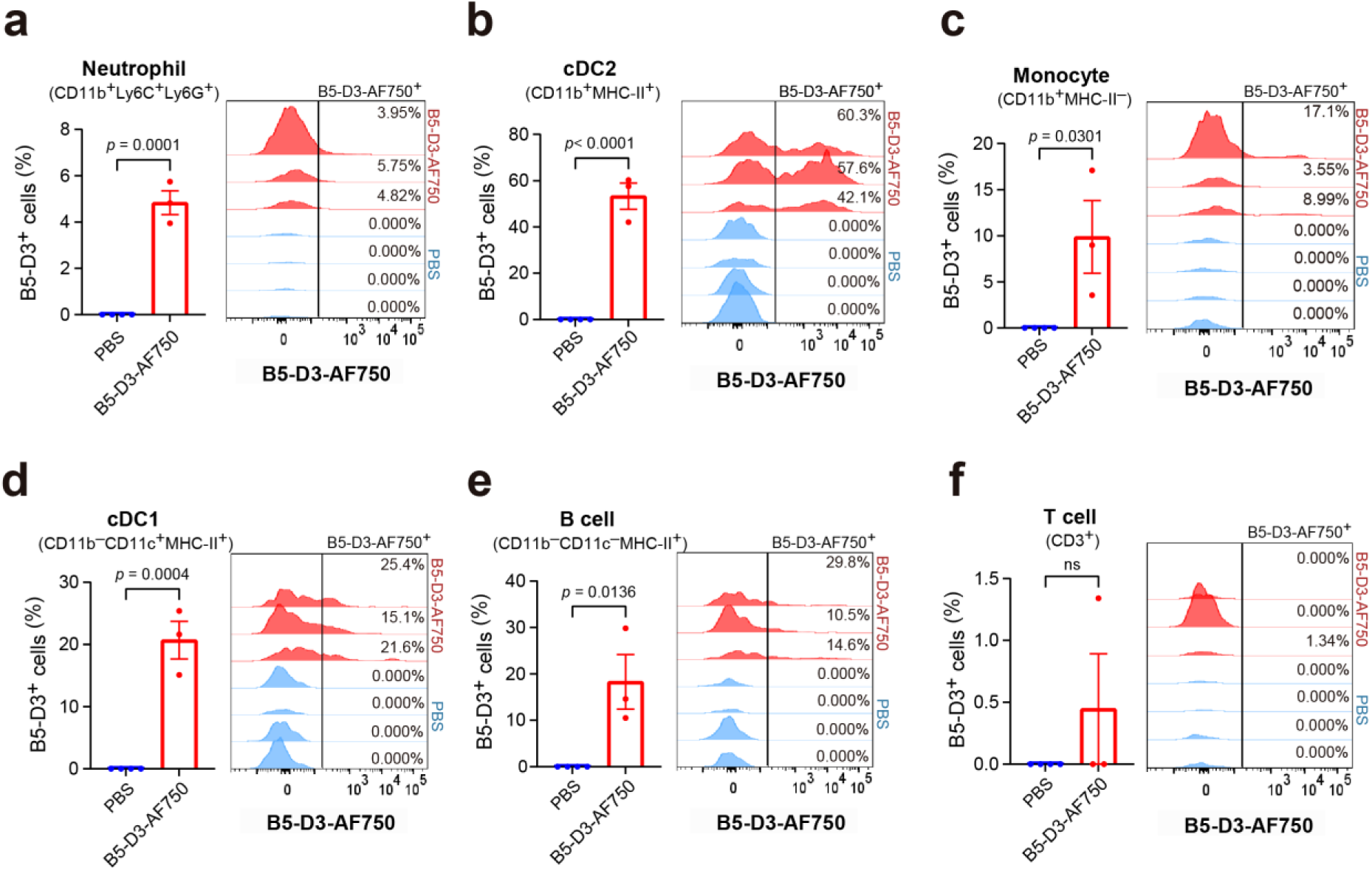
Binding/uptake rates of B5-D3-AF750 in BALF cells. **a**–**f** Positive rates (left) and histograms (right) of B5-D3 binding/uptake as indicated by B5-D3-AF750 fluorescence intensities in CD11b^+^Ly6C^+^Ly6G^+^ neutrophils (**a**), CD11b^+^MHC-Ⅱ^+^ cDC2 (**b**), CD11b^+^MHC-Ⅱ^-^ monocytes (**c**), CD11b^-^CD11c^+^MHC-Ⅱ^+^ cDC1 (**d**), CD11b^-^CD11c^-^MHC-Ⅱ^+^ B cells (**e**), and CD3^+^ T cells (**f**). B5-D3^+^ rates from individual mice are indicated on histograms. Data are presented as mean ± SEM, and statistical significance was determined by Student’s t-test.

**Supplementary Fig. 14.**
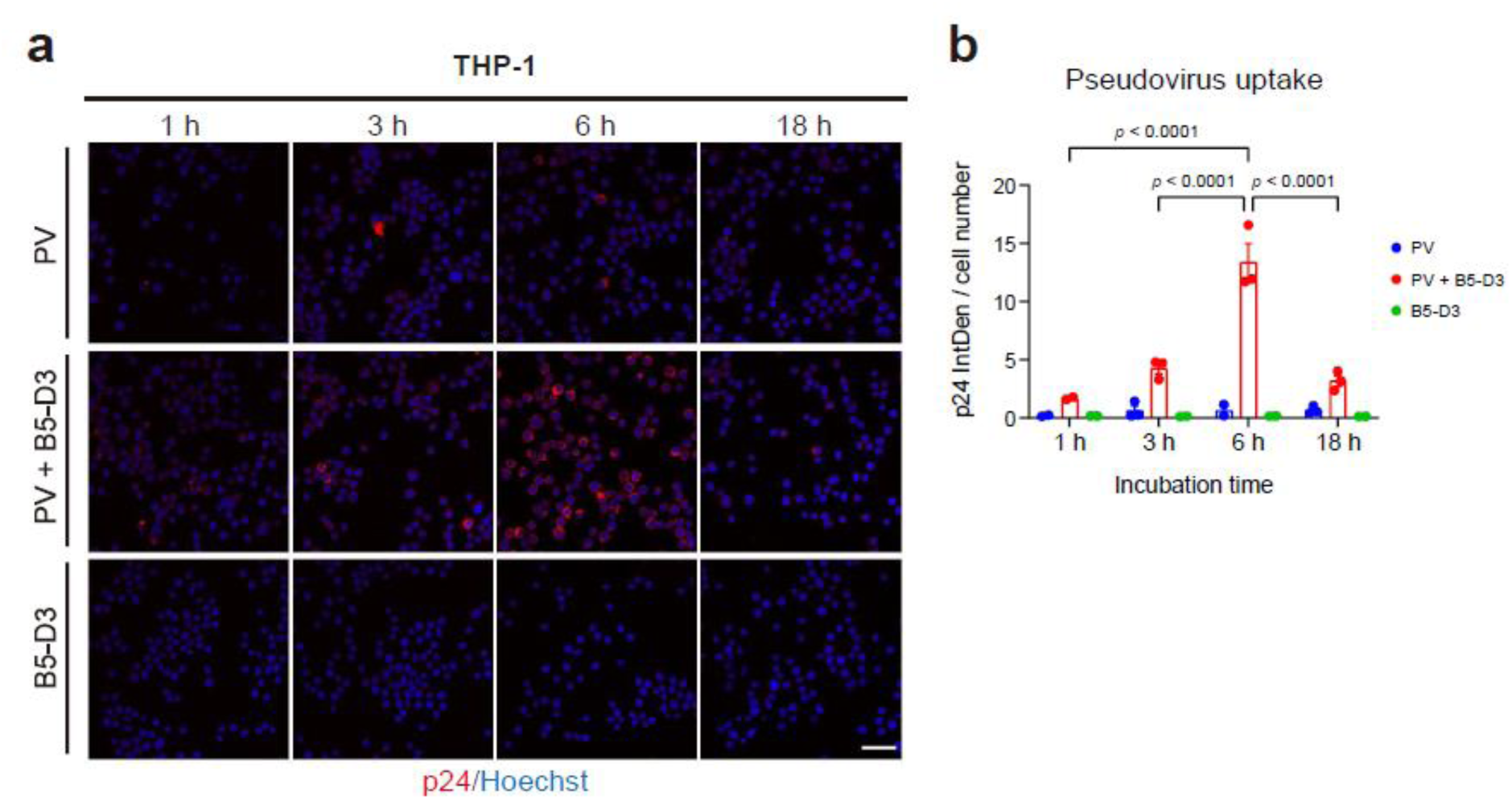
Time-course analysis of sACE2-Fc-dependent pseudovirus entry in THP-1 cells. **a** Representative image showing B5-D3-mediated phagocytosis of pseudovirus by THP-1 monocytes at various time points (1, 3, 6, and 18 h). Cells were incubated with pseudovirus and B5-D3, followed by immunostaining for p24 (red, Invitrogen # PA5-81773). **b** Quantification of mean p24 signal intensity per cell as shown in **a**. IntDen per cell number indicates the average p24 signal per cell, analyzed using ImageJ. Each dot represents one image.

**Supplementary Fig. 15.**
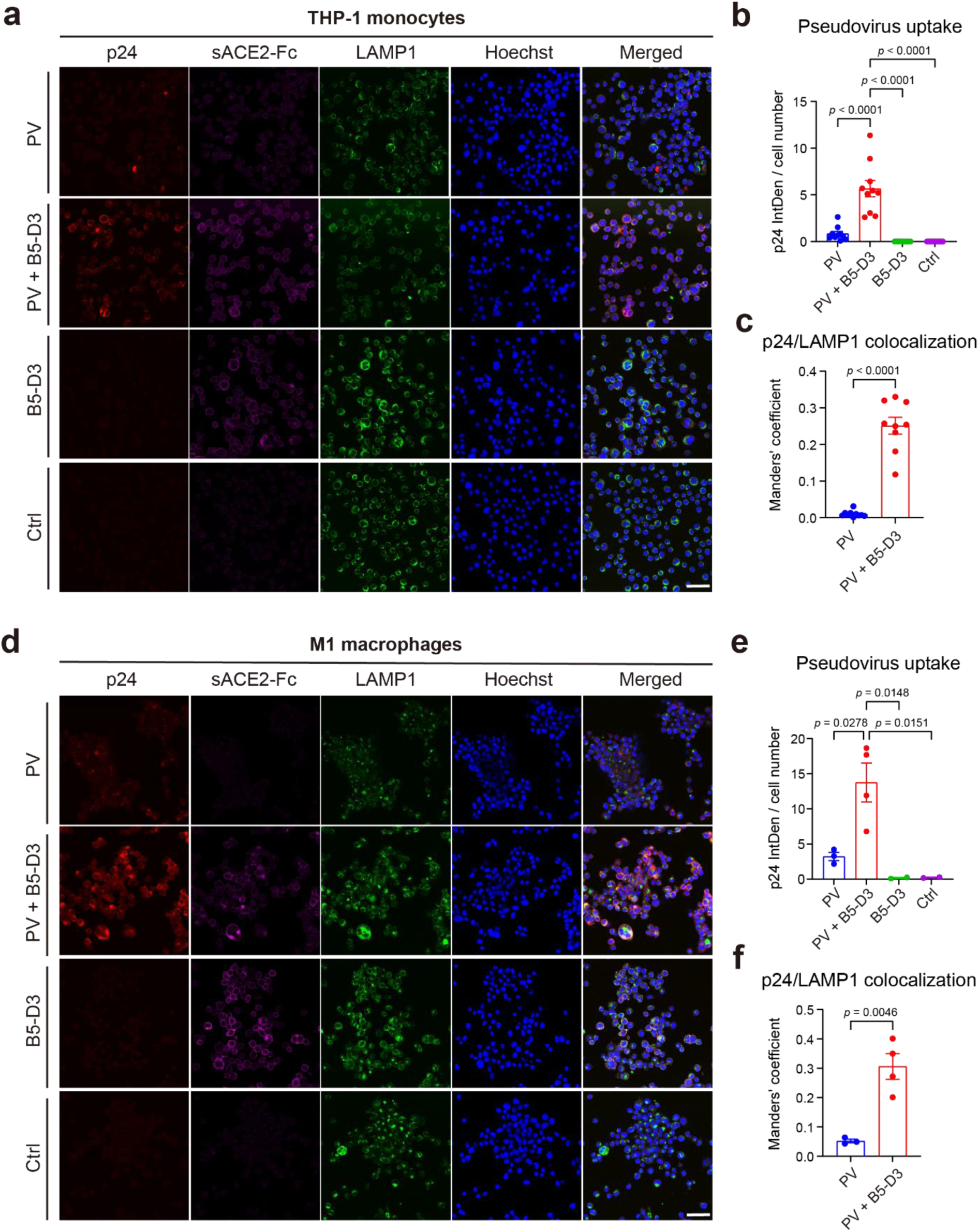
Enhanced phagocytosis of SARS-CoV-2 pseudovirus by THP-1 and THP-1-derived macrophages facilitated by sACE2-Fc. **a**, **d** Representative images illustrating phagocytosis of SARS-CoV-2 pseudovirus by THP-1 monocytes (**a**) and THP-1 differentiated M1 macrophages (**d**) after 6 hours of incubation with or without sACE2-Fc (scale bar = 50 µm). p24 (Invitrogen # PA5-81773) and LAMP1 (Abcam # ab25630) were used to identify the pseudovirus and lysosomes, respectively. **b**, **e** Quantification of mean p24 signal intensity per cell for THP-1 monocytes (**b**) and M1 macrophages (**e**). IntDen per cell number indicates the mean p24 signal per cell, analyzed using ImageJ. Each dot represents one image. **c**, **f** Manders’ coefficient demonstrating the colocalization of p24 and LAMP1 in THP-1 monocytes (**c**) and M1 macrophages (**f**). Data are presented as mean ± SEM, and statistical significance was determined by Tukey’s multiple comparisons test.

**Supplementary Fig. 16.**
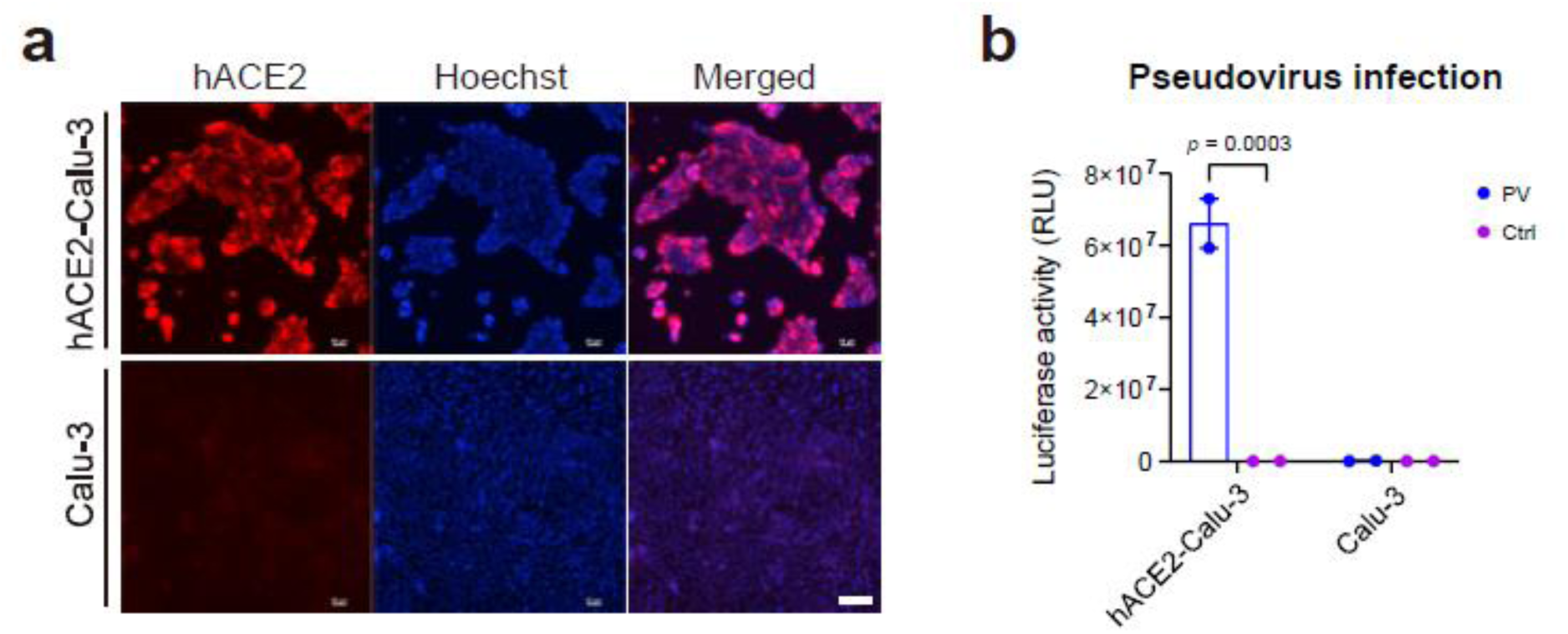
Generation of Calu-3 cell overexpressing hACE2 for enhanced pseudoviral infection. **a** Immunostaining of hACE2 (Abcam # ab15348) in hACE2-Calu-3 and control Calu-3 cells. **b** Infection levels of SARS-CoV-2 pseudovirus in hACE2-Calu-3 versus control Calu-3 cells, quantified by luciferase assay 72 hours post-infection, performed in duplicate. Data are presented as mean ± SEM, and statistical significance was determined by Šídák’s multiple comparisons test.

**Supplementary Fig. 17.**
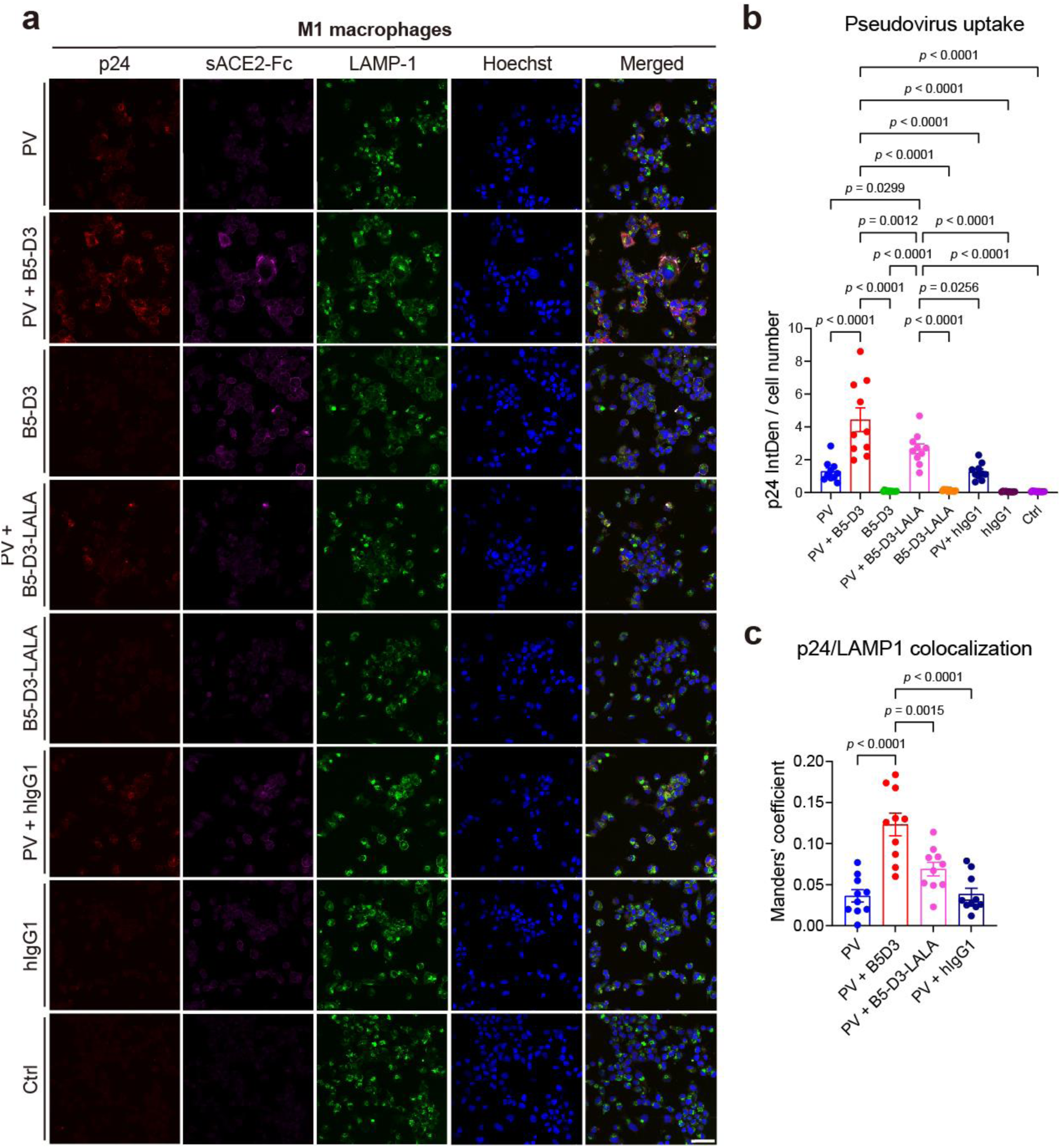
Reduced uptake of SARS-CoV-2 pseudovirus in THP-1-derived macrophages due to malfunction or absence of Fc domain in B5-D3. **a** Representative images illustrating uptake of SARS-CoV-2 pseudovirus (PV) by THP-1-derived M1 macrophages at 6 h after incubation with B5-D3, B5-D3-LALA, and hIgG1 isotype with or without PV. Scale bar = 50 µm. **b** Quantification of mean p24 signal intensity per cell. IntDen per cell number indicates the mean p24 signal per cell, analyzed using ImageJ. Each dot represents one image. Data are presented as mean ± SEM, and statistical significance was determined by Tukey’s multiple comparisons test.

**Supplementary Fig. 18.**
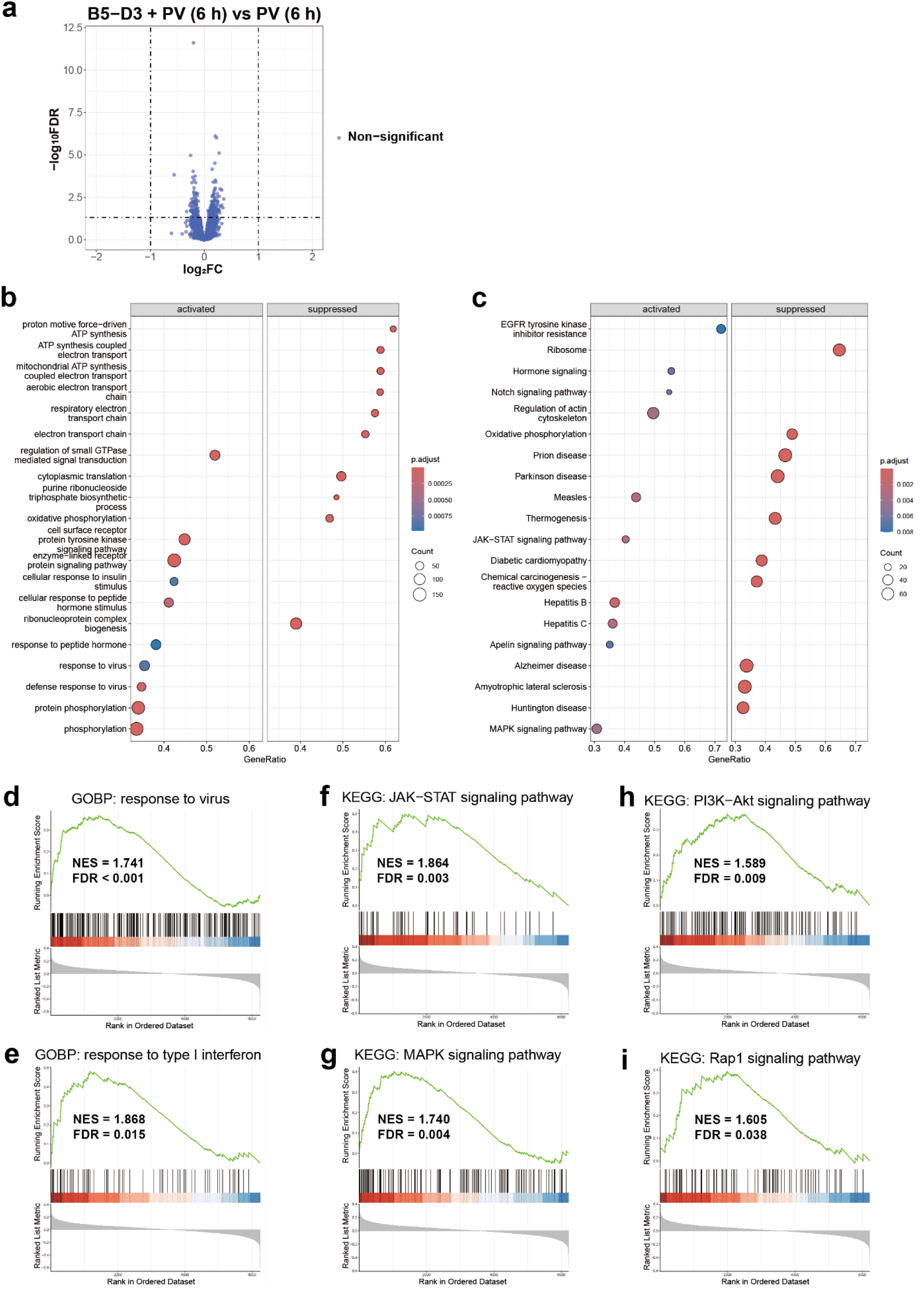
Transcriptomic analysis revealed activation of THP-1-derived macrophages mediated by 6 h incubation with B5-D3-pseudovirus complex. **a** DGE analysis between THP-1-derived M0 macrophages incubated with B5-D3 + pseudovirus (PV) and those incubated with PV only (incubation time = 6 h). **b** GSEA of GOBPs significantly altered in the B5-D3 + PV group compared to the PV group, with top 15 most significantly activated and top 15 most significantly suppressed terms displayed. **c** GSEA of KEGG pathways significantly altered in the B5-D3 + PV group compared to the PV group, with top 15 most significantly activated and top 15 most significantly suppressed terms displayed. **d**–**i** GSEA plots of response to virus (**d**), response to type I interferon (**e**), JAK-STAT signaling pathway (**f**), MAPK signaling pathway (**g**), PI3K-Akt signaling pathway (**h**), and Rap1 signaling pathway (**i**) in B5-D3 + PV vs PV comparison. Benjamin–Hochberg method was used for FDR adjustment.

**Supplementary Table 1.** Enhanced tolerance and stability of B5-derivatives compared to WT sACE2-Fc in AAV-administered K18-hACE2 mice. Detailed *post hoc* comparisons among treatment groups shown in Supplementary Fig. 4b. Diff., difference; **, 0.001 ≤ *p* < 0.01; *, 0.01 ≤ *p* < 0.05. *p* values were determined by Tukey’s multiple comparisons test.

| Tukey's multiple comparisons test | Mean Diff. (ng/mL) | Adjusted *p* | Summary |
| --- | --- | --- | --- |
| PBS vs. WT | -9990 | 0.0017 | ** |
| PBS vs. B5-D3 | -29104 | 0.0046 | ** |
| PBS vs. B5-D4 | -29894 | 0.0028 | ** |
| PBS vs. B5-D5 | -26774 | 0.0013 | ** |
| WT vs. B5-D3 | -19114 | 0.0145 | * |
| WT vs. B5-D4 | -19904 | 0.0078 | ** |
| WT vs. B5-D5 | -16784 | 0.0021 | ** |
| B5-D3 vs. B5-D4 | -790.4 | 0.9459 | ns |
| B5-D3 vs. B5-D5 | 2330 | 0.4915 | ns |
| B5-D4 vs. B5-D5 | 3120 | 0.2103 | ns |

**Supplementary Table 2.**
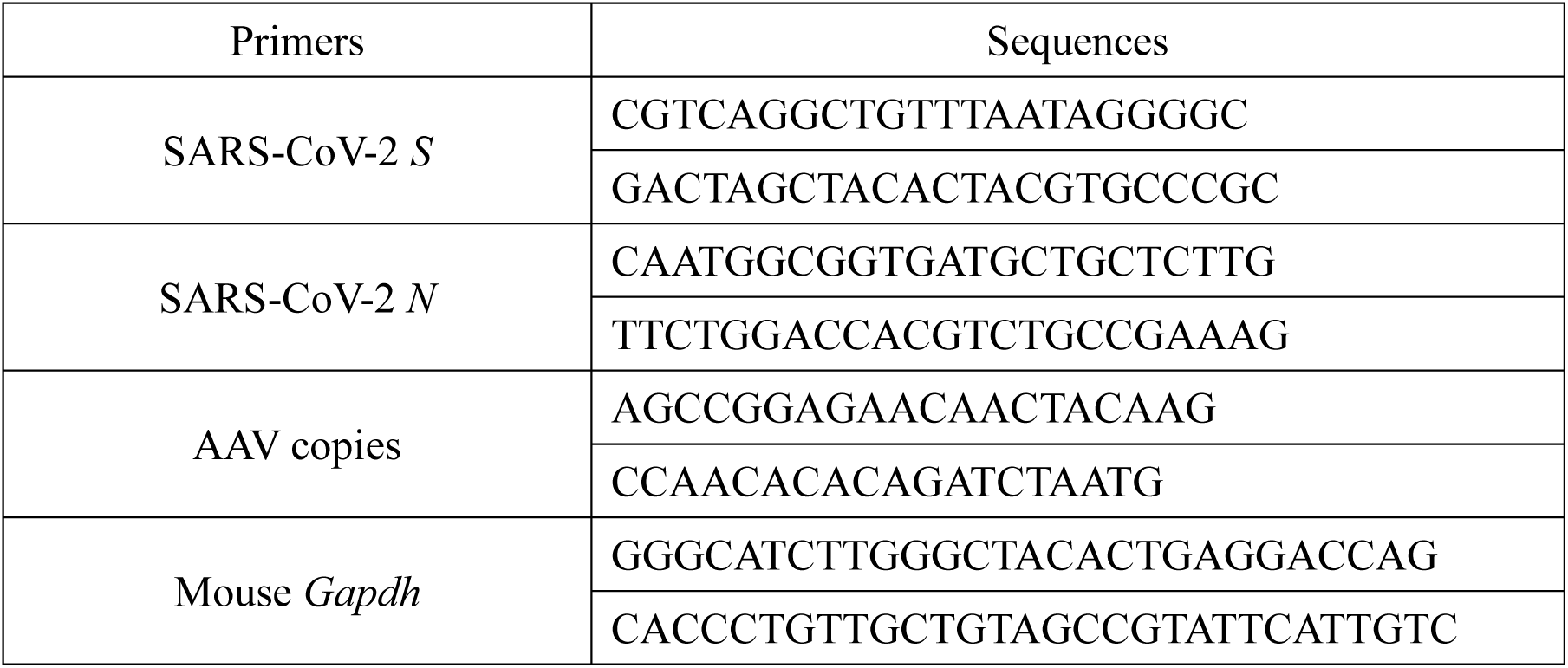
Primers used in quantitative PCR.

